# Plant associated bacteria are a rich reservoir for multidrug efflux pumps

**DOI:** 10.64898/2026.02.24.707749

**Authors:** Liza Rouyer, Yifan Zhu, Martin Parniske, Niklas Schandry

## Abstract

Plants produce antibiotic substances and bacteria need to cope with those substances to colonize plants. We analyzed the inventory of genes encoding resistance-nodulation-cell division (RND) efflux pumps originating from 282 bacterial isolates from leaves or roots of the model plant *Arabidopsis thaliana*. We confirmed that, on average, plant associated bacteria hold a significantly increased repertoire of genes encoding RND antiporters homologs compared to strains isolated from other ecological niches, as reported in a previous study. While some RND antiporter clades were enriched in plant colonizers, other clades found in the genomes of isolates from other environments were underrepresented. An in-depth analysis of RND antiporters from plant colonizing bacteria revealed that conserved motifs can be found in each clade, possibly contributing to substrate specificity. Interestingly, we found horizontal gene transfer markers in 10% of the antiporter homologs, suggesting that horizontal gene transfer may significantly contribute to the adaptation of bacteria to the specific chemical environment created by different organs of plant hosts. In addition, homologs from leaf-isolated bacteria showed a lower diversification, and harbored markers of horizontal gene transfer in the heavy metal exporting clade.

Sequence and structural analysis revealed a high diversity in RND-antiporters, with few residues under purifying selection, indicating that RND diversity is driven by random mutations. Our findings have major implications for the origin of multidrug resistances and for our understanding of the forces shaping the outcome of plant-microbe ecology in general.

## Introduction

Terrestrial plants (hereafter called plants) are a main carbon source for terrestrial microbiota, as they exude up to 40% of their assimilated carbon through the roots (Canarini et al., 2019; Kim et al., 2021; Sasse et al., 2018). Bacteria not only dwell on dead plant material (Drigo et al., 2010) but are also systematically fed by plants that exude organic compounds via their roots (Durán et al., 2018) and leaves (Mercier and Lindow, 2000; Schäfer et al., 2023). These plant metabolites have a dual role, serving as food and consequently growing to high densities on the rhizoplane (Barillot et al., 2013), but often also displaying strain-specific antibiotic activities, thus acting as gate-keepers of the rhizosphere microbiome (Harbort et al., 2020; Schandry et al., 2021). According to our current understanding, the microbiota around the root are systematically maintained by the plant and contribute to plant defense against eukaryotes and pathogens (Bulgarelli et al., 2013).

Plants secure their survival amidst a plethora of multicellular eukaryotic competitors including, fungi, annelids, nematodes, slugs, insects, and vertebrates. All of these organisms rely on plants as their primary nutrient supply. The soil substrate in which most seed plants develop their roots is rich in microbial diversity (Fitzpatrick et al., 2020), with most of these microbes receiving their carbon indirectly via plant photosynthesis. In this microbe-rich environment, plants have evolved chemical strategies to secure their survival. Roots exude a cocktail of chemicals that are thought to nourish and select a specific composition of microbiota that aid the host plant in their battle against pathogenic and/or plant consuming eukaryotes (Parniske, 1991; Parniske et al., 1991; Sasse et al., 2018; Voges et al., 2019).

To colonize the rhizosphere environment, bacteria need to be able to deal with the chemical stress imposed by the plant. One strategy employed by bacteria to defend against antibacterial compounds is to export the toxic compound through multidrug-resistance pumps, as known from tetracycline resistance (Blanco et al., 2016; Chopra and Roberts, 2001). Among the five major families of multidrug-resistance pumps, Resistance-Nodulation-cell Division (RND) antiporters have received particular attention because of their contribution to multi-drug-resistances in pathogenic bacteria (Blair et al., 2014). RND transporters are tripartite protein complexes, consisting of the inner membrane localized, periplasmic, substrate specificity conferring homo-trimeric antiporter, which is connected to the channel via an adaptor protein. The model RND transporter in *E. coli* is the AcrAB-TolC transporter, consisting of the antiporter AcrB, the adaptor AcrA, and the channel TolC. We will use the terms *antiporter*, *adaptor*, and *channel* throughout the text to refer to the RND components homologs to *acrB, acrA*, and *tolC*, respectively. Because of their clinical relevance, it is perhaps not surprising that the structure of the specificity conferring antiporter component AcrB was already resolved over 20 years ago (Murakami et al., 2002; Yu et al., 2003). Accumulating evidence shows that RND-type efflux pumps play important roles in Gram-negative bacterial survival and virulence (Fanelli et al., 2023; Wang-Kan et al., 2017; Zahedi Bialvaei et al., 2021). For example, the TtgABC efflux pump, a member of the RND superfamily, has been experimentally confirmed to export multiple antibiotics, including chloramphenicol and tetracyclines, and is regulated in an antibiotic-dependent manner by the TetR-family transcriptional regulator TtgR in *Pseudomonas putida* (Terán et al., 2003)

Bodilis *et al*. (2024) asked whether there are any relationships between the number and categories of RND-antiporters in a bacterial genome, and the ecosystem they originate from. From 920 representative genomes of Gram-negative bacteria isolated from nine different environments, they identified 6205 genes encoding RND-antiporters with an average of 6.7 genes per genome. Remarkably, they observed an enrichment of HAE-1 subfamily RND-antiporter genes in 67 genomes from plant-isolated bacteria. They further validated that rhizosphere-isolates bacteria from the plant *Fallopia japonica* possess significantly higher numbers of the HAE-1 subfamily antiporter than adjacent bulk soil. Recently, an inspection of the RND repertoire of 185 Arabidopsis root-associated bacterial genomes found on average 11 RND homologs per genome (Russ et al., 2025). Importantly, they identified one RND encoding operon, which includes the gene encoding the ef90 antiporter, conferring resistance to toxic, plant-derived glucosinolate breakdown products, indicating a host plant specific functional adaptation (Russ et al., 2025).

Using a collection of 282 Arabidopsis-associated Gram-negative bacterial genomes (Bai et al., 2015), we asked to what extend the observations by Bodilis and colleagues (Bodilis et al., 2024) regarding the elevated number of RND-antiporter homologs in the root environment can be confirmed for *Arabidopsis thaliana*. By comparing the distribution of RND-antiporter homologs in root- and leaf-associated bacteria, we investigated whether the enrichment of homologs in plant-associated bacteria is specific to the rhizosphere compartment or extends to the phyllosphere. Moreover, we asked whether the *ef90* antiporter from North Carolina soil (Russ et al., 2025) could also be found in a collection of bacterial isolates from Arabidopsis sampled across five locations in Germany and France (Bai et al., 2015). We investigated the prevalence of horizontal gene transfer and found an enrichment of HGT events in leaf-associated bacteria. Sequence and structure conservation analysis across all clades showed a lack of conservation in the substrate export domains, which could be further confirmed by the absence of conserved motifs in the substrate entry channel.

## Material and methods

### Bacterial genomes and taxonomy

The genomes of 282 Gram-negative bacteria were downloaded from the AtSphere database (Bai et al., 2015) and annotated using Bakta (Schwengers et al., 2021) to obtain the translated protein sequences. To build the phylogenetic tree of Figure 1A, we BLAST’ed (v2.5.0+, Camacho et al., (2009)) the predicted proteomes against the amino acid sequences of five conserved genes (atpD (NCBI ID: CAD6023125.1), dnaJ (NCBI ID: NP_414556.1), gyrB (NCBI ID: CAD5998225.1), recA (NCBI ID: CAD6005917.1) and rpoB (NCBI ID: NP_418414.1)). The alignment of all homologs was used to build a neighbor-joining phylogenetic tree.

**Figure 1.**
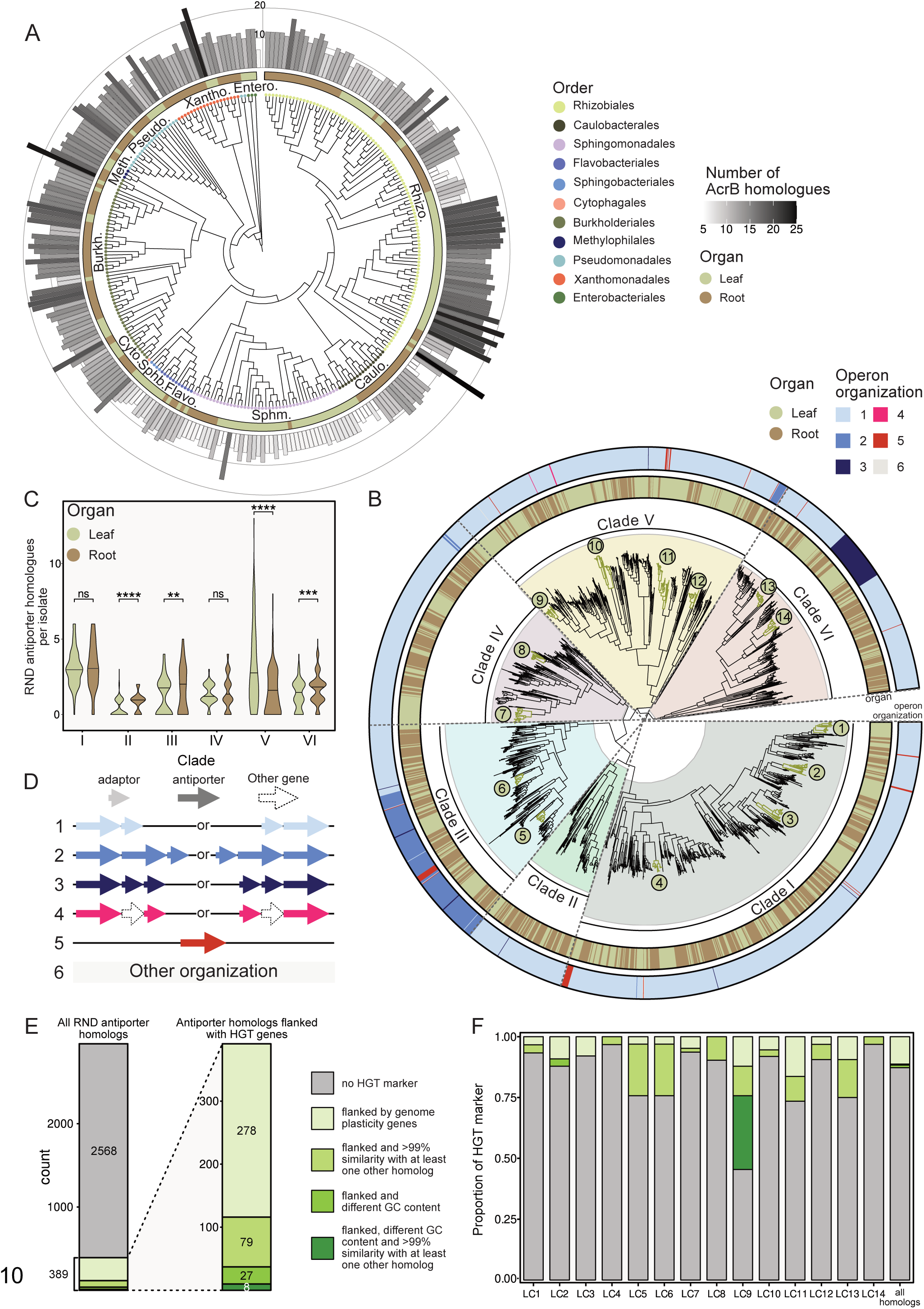
Plant-associated bacteria harbour multiple copies of highly diversified RND export pumps. **A.** Phylogenetic tree of 282 Gram-negative bacterial isolates, based on the amino acid sequence alignments of five conserved proteins (See M&M Bacterial genomes and taxonomy). Inner nodes show taxonomical order, the outer ring the organ (roots or leaves) of *Arabidopsis thaliana* from which they were isolated. Bar plot height and shade displays the number of RND-antiporter homologs identified in each genome. **B.** Protein sequence similarity tree of RND export pumps (see M&M Homology search and phylogeny). Inner ring: origin (root or leaf) of the isolate carrying each RND-antiporter homolog. Green numbered dots highlight clusters of 30 or more closely related homologs from genomes of different bacteria isolated from the same organ. Outer ring: operon organization for each homolog. **C.** Number of encoded RND-antiporter homologs per isolate separated by clade and root or leaf origin of the isolate. p-values obtained with a Wilcoxon signed rank test. **: p<0.01, ***: p<0.001, ****: p<0.0001, ns: not significant. **D.** Different RND operon organizations shown in panel B. **E.** Number of homologs displaying different type of horizontal gene transfer (HGT) markers. **F.** Proportion of HGT markers in all leaf clusters (LC) highlighted in panel B and in all homologs of this study.

### Homology search and phylogeny

An initial set of RND antiporter homologs was detected by performing a BLASTp (protein-protein BLAST v2.5.0) search on the bacterial protein sequences using an *E. coli* AcrB sequence as reference (NCBI ID: NP_414995.1). The sequences obtained were filtered for e-value <10E-4 and length >600aa. The resulting 1966 sequences were aligned with MAFFT with custom parameters (--op 1, --ep 0.1, Katoh et al., (2002)) via the Nextflow pipeline nf-core/multiplesequencealign (Santus et al., 2024, Katho et al., 2002). The alignment was used to build a Hidden-Markow-Model profile using HMMER (v3.4, hmmer.org), which was subsequently used for searching against the bacterial genome database with default parameters. The obtained sequences were filtered for an e-value < 10e-4, length > 600aa, and HMM accuracy > 0.85, resulting in a total of 2960 putative RND antiporter homologs. The same method was used to find periplasmic adaptor protein homologs, starting for the *E. coli* AcrA protein (NCBI ID: NP_414996.1) without length filter. Only putative homologs that were localized on the same strand as AcrB homologs and within up to 5 genes up/downstream were kept.

To build the phylogenetic tree, the sequences of all homologs were aligned with MAFFT using the same parameters as above. Maximum likelihood phylogeny trees were built using IQTREE2 (v2.3.6; Hoang et al., 2018; Kalyaanamoorthy et al., 2017; Minh et al., 2020) with 1000 ultrafast bootstraps. The best fitting model (LG+F+R5) according to the Bayesian Information Criterion was determined using 10 individual random set of 50 sequences and applied for the building of the complete tree (Figure 1B). The tree was rooted with 58 homologs that did not form a defined monophyletic clade (Supplementary Figure 1B).

### Comparison to previous studies

To compare the 2960 RND antiporter from this study to previous studies, we incorporated 83 characterized RND antiporters from the TCDB database (full list in Supplementary Table S5; the same templates used in Bodilis et al., 2024; Saier et al., 2006) for the phylogenetic analysis (Supplementary Figure S1A). Multisequence alignment of the 3043 amino acid sequences was performed with MAFFT (same parameters as above), followed by maximum-likelihood phylogenetic tree construction with IQ-TREE2 (v2.3.6; 1000 ultrafast bootstrap replicates). The best fitted model (LG+F+R10) was selected, and the tree was rooted with a monophyletic clade containing a subset of the outgroup in Figure 1B. The resulting tree was visualized in R (v4.3.1) using ggtree (Yu, 2020) and ggtreeExtra (Xu et al., 2021) packages (Supplementary Figure S1A).

The list of RND antiporters and their bacterial origins was retrieved from Supplementary Table 1 of Bodilis et al. (Bodilis et al., 2024). Bacterial origins represented by fewer than four isolates or assigned to multiple environment categories were excluded prior to analysis to minimize potential bias. To distinguish between plant-associated and non-plant-associated bacteria, we recalculated the average number of RND antiporters per isolate for each clade defined by Bodilis et al., (2024) in their study. Statistical comparisons were performed using the Wilcoxon signed-rank test (Supplementary Table S2).

To investigate the distribution of the *ef90* operon identified in Pseudomonas strains from North Carolina Arabidopsis fields (Russ et al., 2025), we retrieved the PS417_13190 protein sequence (GenBank accession AIB36522.1) and used it as the query for a BLASTp search against 2960 RND antiporter homologs identified from 282 Arabidopsis-associated genomes, as well as 83 homologs obtained from the TCDB database. Top-hit homologs (21 sequences) in our dataset with over 80% sequence similarity to the query are labelled in Supplementary Figure 1A.

### Consensus sequence, structure prediction and analysis

Consensus sequences and conservation values (Figure 2A) were obtained from the MAFFT alignments using the R package Bio3D (Grant et al., 2006). The resulting sequences were used as an input for the AlphaFold server (AlphaFold3, Abramson et al., 2024). The predictions with the highest confidence scores were used for subsequent analysis. The protein domains were determined by aligning our consensus sequences to well characterized RND transporters as reference (Kobylka et al., 2020; Murakami et al., 2002). RMSD values were calculated with USCF ChimeraX (v1.8; Meng et al., 2023) using the Matchmaking tool and the predicted structure of the consensus sequence of all RND transporter homologs found in this study as reference (Figure 2C).

**Figure 2.**
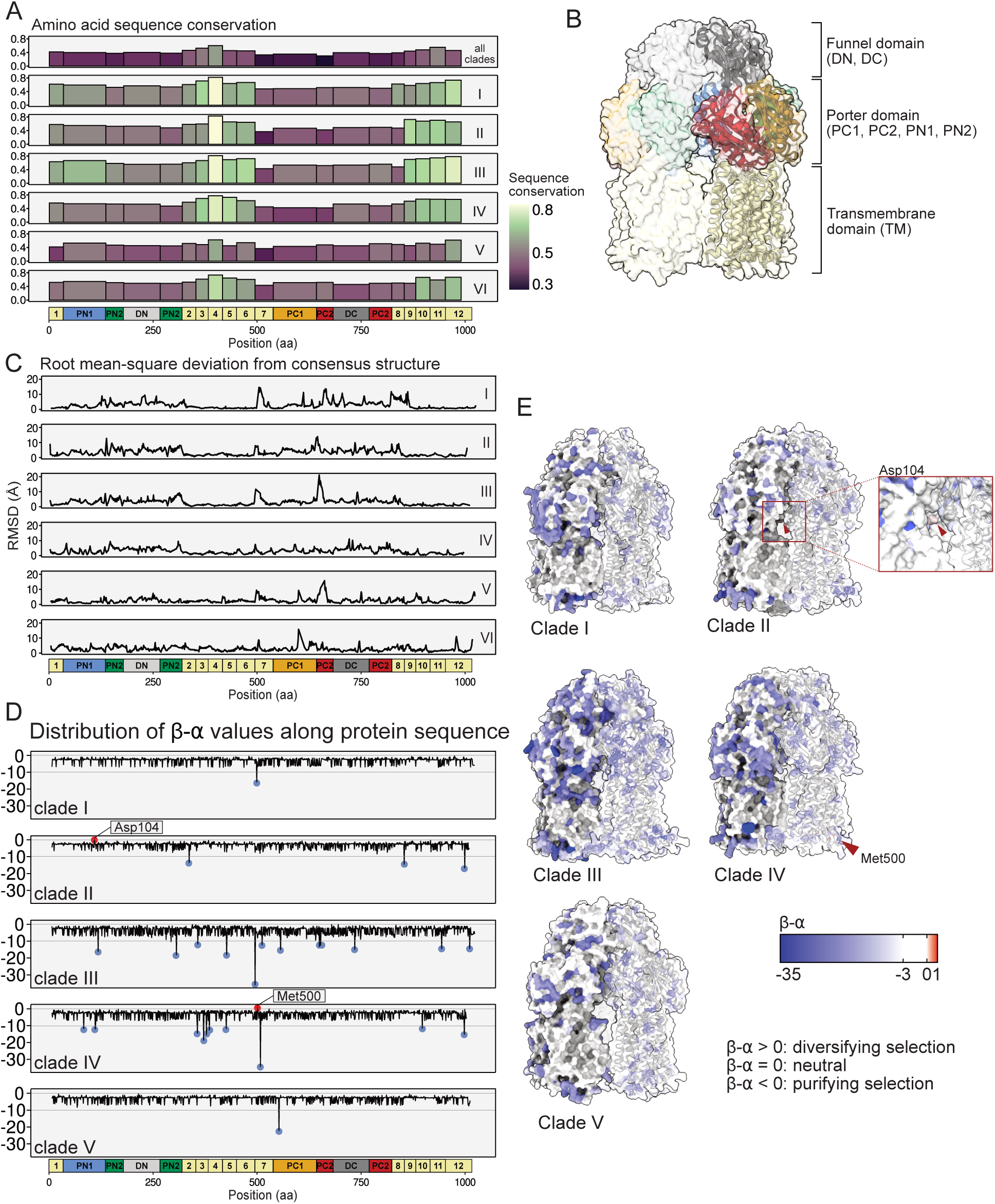
Sequence and structure conservation differs by domain. **A.** Mean amino acid sequence conservation of RND-antiporter per protein domain across all clades (top) and individual clades. Bar height and colour shows sequence conservation. Conservation per residue can be found in Supplementary Figure S3. **B.** Predicted trimeric protein structure of the consensus sequence of all RND-antiporters, coloured by domains, lateral view. **C.** Root-mean-square deviation (RMSD) of atomic positions of protein structures predicted from consensus sequences, comparing consensus structures of individual clades to the consensus structure of all homologs (shown in A). **D.** Distribution of β-α values along the amino acid sequence. Positive values are indicated in red, values below -10 are indicated in blue. **E.** Projection of the β-α values on the predicted structure of the consensus sequence from each clade. Residues under diversifying selection are highlighted in red and indicated with a red arrow. Protein structure predictions were done with AlphaFold3 on the AlphaFold Server. See M&M “Consensus sequences, structure prediction and analysis” for more details.

### Selection inference

Selection inference on each residue (Figure 2D) was performed using FUBAR (Murrell et al., 2013) as part of the HYPHY software (v2.5.29; Pond et al., 2005). As FUBAR requires codon-aware alignments, the sequences were pre-processed following the codon-msa scripts from https://github.com/veg/hyphy-analyses. Briefly, the coding sequences for each homolog was corrected for frame-shift mutations and translated to amino acids (default parameters for clades A, B and C, custom E number (fraction of N allowed in the alignment) for clade D (0.01) and clade E (0.005). The obtained amino acid sequences were aligned using MAFFT (same parameters as above). The protein multiple sequence alignment (MSA) and the frameshift corrected nucleotides sequences were then used to obtain a nucleotide MSA.

The nucleotide MSA was used to build a phylogenetic tree using IQTREE2 (see above, best model according to ModelFinder GTR+F+I+R10), used as an input for FUBAR along with the nucleotide MSA. The obtained files were imported into HyPhy Vision for export to tables and subsequent analysis in R.

### Motif discovery

Motif discovery (Figure 3) was performed on all homologs and on each clade separately with STREME (v1.16.0; Bailey, 2021). For the analysis with all homologs, the outgroup was used as negative control. For the clade-specific analysis, the sequences of all other clades were used as negative control.

**Figure 3.**
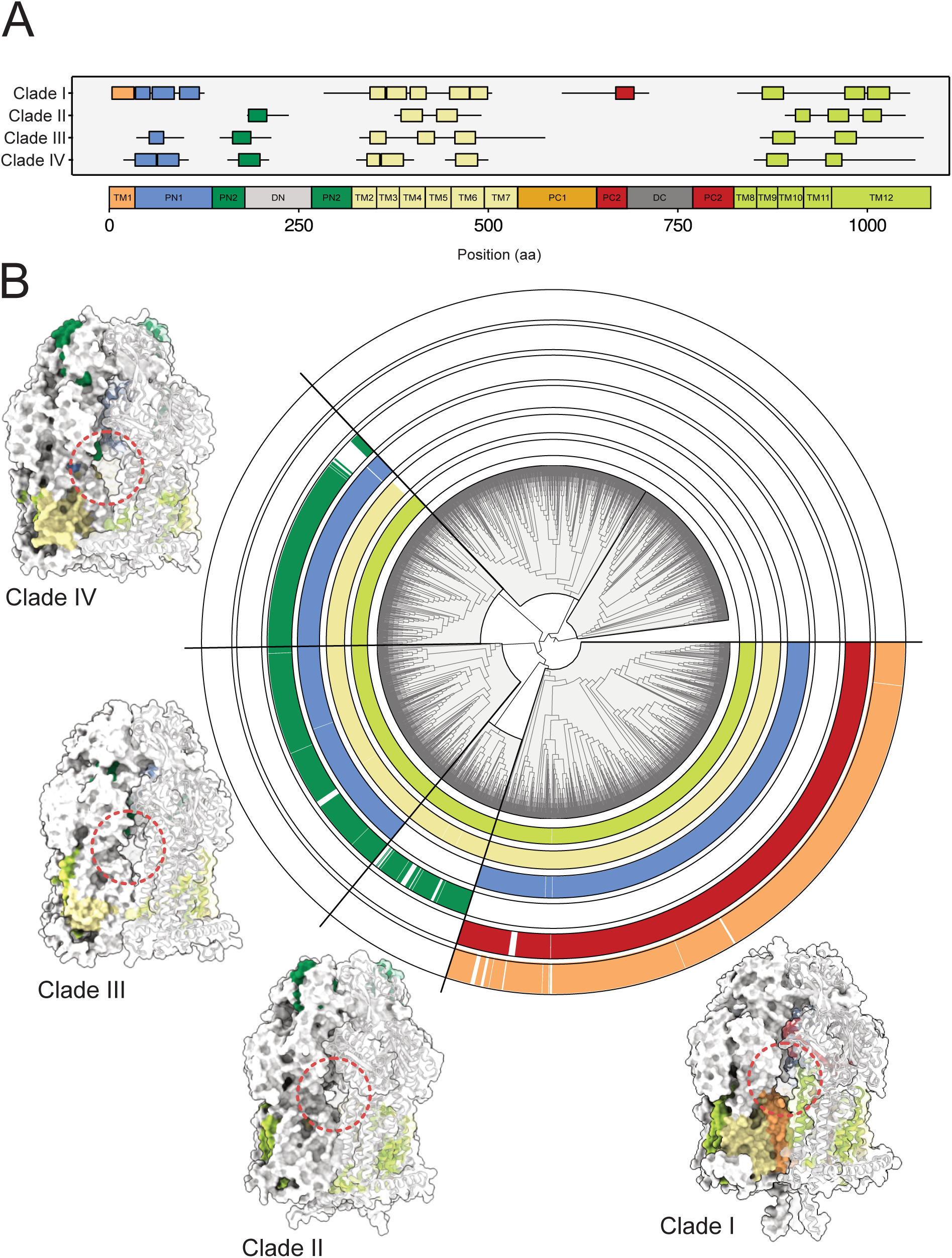
RND transporters harbour clade-specific conserved motifs. **A.** Sequence position of clade-specific motifs present in >75% of homologs of each respective clade. Clades V and VI do not have conserved motifs. The box indicates the mean start and end of each motif across all homologs of a given clade, whiskers show the distribution of the start and end of the motifs across the homologs. **B.** Distribution of the conserved motifs on the structure-similarity tree of all RND transporter homologs and localization of the conserved motifs on the predicted protein structures of selected representative of each clade. Branch length is not proportional to taxonomic distance. Red dotted circles indicate the putative substrate entry channels based on known RND transporter structures. See M&M Motif discovery for details.

### Statistical analysis and data visualization

All statistical analysis (operon organization, organ of origin, general statistics, clade determination) and data visualizations were carried out on R (v4.4.2) using custom scripts available on https://gitlab.plantmicrobe.de/nschandry/rnd-transporters-in-plant-associated-bacteria.

## Results

### Plant associated bacteria harbor multiple copies of highly diversified RND antiporters

We analyzed the genomes of a total of 282 bacteria of which 148 were isolated from the rhizosphere and 134 from the phyllosphere of *Arabidopsis thaliana* individuals, collected from five soils (rhizosphere) and six sampling sites (phyllosphere) (Supplementary Table S1, Bai et al., 2015). The phylogeny of the isolates, based on the amino acid sequences of five conserved proteins (see M&M Bacterial genomes and taxonomy for details), showed a diverse representation of 11 bacterial orders, each comprising leaf and root isolates (Figure 1A, Supplementary Table S1). Using a hidden-Markov-model based sequence search on the newly annotated genomes, we identified a total number of 2960 RND-antiporter homologs in the genomes of all isolates. Individual genomes harbored between 1 and 26 RND-antiporter homologs, with an overall average of 10.5 homologs per genome (Figure 1A, Supplementary Table S1). This number was only marginally different between isolates originating from root or leaves.

Our analysis revealed a higher average number of RND antiporters per genome than reported by Bodilis and colleagues, who found an average of 6.7 RND homologs per genome (Bodilis et al., 2024). Because this dataset included genomes of bacteria from multiple origins, we retrieved the RND antiporter list for all 920 genomes in their study and reanalyzed data to enable a direct comparison of RND antiporter copy numbers between plant and non-plant-origin genomes (see M&M Comparison to previous study). Among the 920 genomes from nine environments, the 67 plant-origin genomes harbored an average of 10.9 RND antiporter homologs per genome, representing a 1.8-fold enrichment relative to genomes from all other environments (Supplementary Table S2), which is consistent with our results (Figure 1A).

We examined the RND-antiporter content across plant-colonizing bacterial lineages and observed significant variation between isolates. On average, isolates from the orders Methylophilales, Burkholderiales, Xanthomonadales, Pseudomonadales, Rhizobiales, and Enterobacteriales possess more than 10 RND antiporter homologs per genome. In contrast, Sphingomonadales carry fewer RND antiporter homologs, with an average of 6.3 copies per genome (Supplementary Table S3). In particular, within the order Rhizobiales, species of the family Hyphomicrobiaceae possess an average of only 3.4 RND antiporter homologs per genome, while Methylobacteriaceae from the same order harbor an average of 15.3 antiporter homolog per genome (Supplementary Table S3).

### RND antiporters from plant-associated bacteria fall into six monophyletic clades

We built a phylogenetic tree based on predicted protein sequences of all 2960 RND-antiporter homologs (see M&M Homology search and phylogeny for details). This analysis revealed six monophyletic clades (Figure 1B, Supplementary Figure S1A). All clades were supported by high bootstrap values (100, 100, 100, 95, 74, 89 for Clades I, II, III, IV, V, and VI, respectively, Supplementary Figure S1B). To analyze whether and to what extend these phylogenetic boundaries correlate with different exported substrates, we included previously functionally characterized RND-antiporters available on Transporter Classification Database (TCDB) from multiple independent sources, aligned the amino acid sequences with the ones from our dataset, and reconstructed the phylogenetic tree (see M&M Comparison to previous study). We labelled the substrates of the functionally characterized RND antiporters in the phylogenetic tree and found that the different families of RND-antiporters cluster in distinct clades (Supplementary Figure S1A, Table 1). Similar to the results of Bodilis and colleagues (Bodilis et al., 2024), members of the heavy metal exporters (HME) family (Saier Jr et al., 1994) all clustered in one single clade (Clade V in our case). Clade IV contains homologs of the nodulation factor exporter (NFE) family (Saier Jr et al., 1994), and Clades I, II, III and VI contain homologs of the larger hydrophobe/amphiphile efflux (HAE) multidrug export family (Saier Jr et al., 1994). The single ancestral duplication reported in clade *B* by Bodilis et al. (2024) can also be seen in Clade III (Supplementary Figure S1B, Table 1).

**Table 1.**
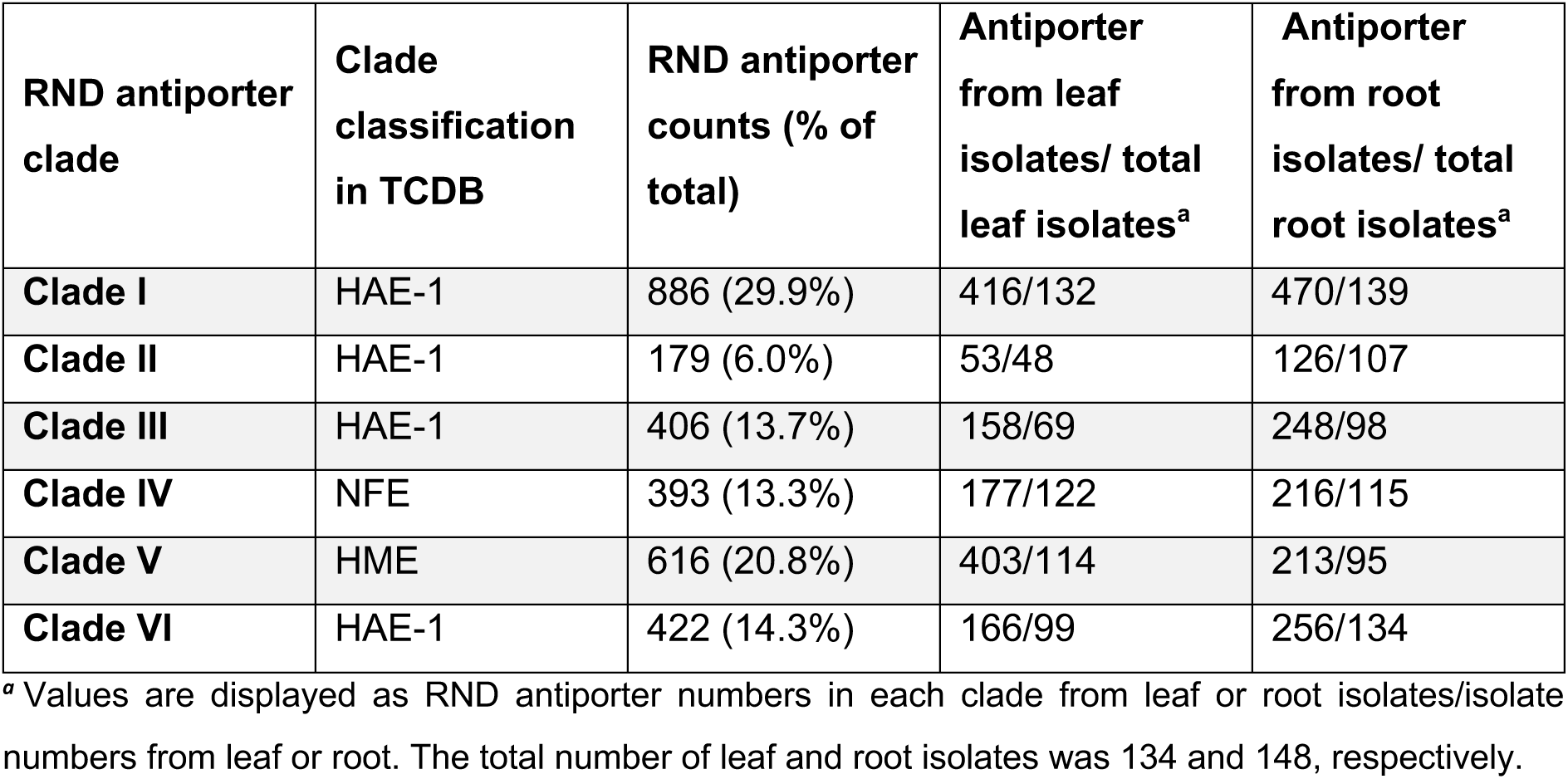
Distribution of antiporter homologs across clades and organs.

| RND antiporter clade | Clade classification in TCDB | RND antiporter counts (% of total) | Antiporter from leaf isolates/ total leaf isolates<sup>a</sup> | Antiporter from root isolates/ total root isolates<sup>a</sup> |
| --- | --- | --- | --- | --- |
| **Clade I** | HAE-1 | 886 (29.9%) | 416/132 | 470/139 |
| **Clade II** | HAE-1 | 179 (6.0%) | 53/48 | 126/107 |
| **Clade III** | HAE-1 | 406 (13.7%) | 158/69 | 248/98 |
| **Clade IV** | NFE | 393 (13.3%) | 177/122 | 216/115 |
| **Clade V** | HME | 616 (20.8%) | 403/114 | 213/95 |
| **Clade VI** | HAE-1 | 422 (14.3%) | 166/99 | 256/134 |

<sup>a</sup> Values are displayed as RND antiporter numbers in each clade from leaf or root isolates/isolate numbers from leaf or root. The total number of leaf and root isolates was 134 and 148, respectively.

Of the nine monophyletic clades defined by Bodilis et al., seven contained significantly higher copy numbers of RND antiporters in plant origin isolates, with the 5-fold enrichment observed in clade *B* (Supplementary Table S2). In our dataset, clades *C*, *D*, and *E* of Bodilis et al. formed a single monophyletic Clade IV (bootstrap support 95%), whereas clades *F*, *G*, and *I* grouped within Clade VI (bootstrap support 89%; Supplementary Figure S1).

The relative distribution of homologs across all the clades is similar to what was reported in Bodilis et al., with Clades I, II, III and VI (HAE) together comprising 63.9% of the sequences (Table 1), highlighting the wide distribution of members of the HAE clades among plant-associated bacteria. While all bacterial orders harboured homologs from each clade, some differences could be observed at the isolate level (Supplementary Figure S2). Only few Sphingomonadales genomes contained homologs from Clades II and III (3/43 and 1/43, respectively), while Pseudomonas genomes generally lacked homologs from Clade IV (24/25) (Supplementary Figure S2). Interestingly, over half (65/105) of the Rhizobiales genomes had no homologs from Clade V, which corresponded largely to the root-associated isolates. Indeed, genomes of 18/52 leaf isolates and 47/53 root isolates did not have homologs from Clade V (Supplementary Figure S2), suggesting a difference in the heavy-metal exposure or detoxification of Rhizobiales depending on their organ of isolation.

We also found 21 homologs with high (over 84%) sequence similarity to the *ef90* RND antiporter identified by Russ et al. (2025) clustering within one subclade of our clade I (Supplementary Figure S1A), suggesting that *ef90*-like antiporters with potential detoxifying functions are widespread in the Cologne soil communities studied.

### Remarkable expansion of the nodulation factor exporter (NFE) clade in plant-associated orders of Rhizobiales and Burkholderiales

One notable difference to previous findings is that the NFE clade is distributed throughout our clade IV, with the majority of sequences clustering in a small subclade (Supplementary Figure S1A). In our dataset, this small subclade, which contains the original NolG of *Sinorhizhobium meliloti* (TCDB ID: 2.A.6.3.1), represents 2.6% (76/2960) of sequences, about 3-fold larger than the proportion reported by Bodilis et al. (0.8% of 6920 sequences). If the entire clade IV is considered according to NolG-homology distribution, it accounts for 13.3% (393/2960) of sequences in our sample set, a 16-fold increase compared to previous studies (Bodilis et al., 2024). The original NolG was identified and named due to its slight root hair deformation defective phenotype and its physical proximity to the cluster of common nod genes of *S. melilotii* (Baev et al., 1991), which are required for Lipo-chitooligosaccharide (LCO) synthesis (Truchet et al., 1991). To the best of our knowledge, all later reported members of the NFE clade were based on the shared global homology with other RND-antiporters without experimental support for their function in LCOs transportation (Barnett et al., 2001; Black et al., 2012). Interestingly, putative symbiotic Rhizobiales account for less than half of clade IV (41%, Supplementary Table S4). Antiporters from other bacterial orders, particularly Sphingomonadales, Caulobacterales, and non-symbiotic members, also contribute substantially to this clade (Supplementary Table S4).

### Leaf and root-specific distribution of phylogenetic clades of RND transporters

We compared the quantitative representation of all RND-antiporter clades in a total of 134 leaf and 148 root isolates (Figure 1B, 1C and Table 1). All clades contain RND-antiporter homologs from genomes of bacteria isolated from leaves and roots, indicating that the organ of origin does not lead to significant phylogenetic clustering of the RND-antiporter repertoire (Figure 1B, Supplementary Figure S2). However, antiporters from leaf isolates are overrepresented in Clades II and V, with on average each leaf-isolated bacterium harboring 0.85 and 3 homologs from these clades, respectively, versus 0.4 and 1.4 homologs per root-isolated bacterium (Figure 1B, C, Table 1). Conversely, antiporters from root isolates are more abundant in Clades III and VI, with 1.7 versus 1.2 homologs per leaf- and root-isolated bacteria on average for both clades (Figure 1B, C and Table 1).

### Quantitative distribution of different operon structures across clades

Analysis of the RND-antiporter-containing operon organization revealed 5 major operon variants (Figure 1B, D). The absolutely conserved feature across variants 1, 2 and 3 (94%; 3032/3239) is an RND-antiporter homolog localized in the immediate vicinity of an RND-adaptor homolog, with differences in the ratios between antiporter and adaptor genes in the different operon variants. Clade III holds a total of 406 RND-antiporters, of which 118 are arranged in variant 1 (one antiporter, one adaptor), while 273 are arranged in variant 2, a tandem arrangement of two RND-antiporters immediately next to an RND-adaptor, suggesting gene duplication or gene insertion events. Similarly, 89 homologs (20%) from Clade VI showed a pattern of two tandemly arranged RND-adaptors paralogs immediately 5’ of one RND-antiporter homolog (variant 3). We tested to what extend this tandem array could be the consequence of recent and local duplication events in this clade, and found that the homologs in close proximity to each other had between 52% and 77% identity to each other and to other homologs within the same genome. This suggests that the tandem arrangement is not due to a recent local duplication, but could be the cause of a more ancient local duplication or genome rearrangement.

### Horizontal gene transfer in RND antiporter homologs

We investigated the prevalence of horizontal gene transfer (HGT) events among all RND antiporter homologs. Signatures of HGT across bacterial genomes include: high sequence identities between the sequences subject to HGT, flanking regions involved in mobilisation, and differences in GC content between the transferred sequence and the genomic background (Arnold et al., 2022). We identified 389 homologs that harbored flanking genes related to genome plasticity and gene transfer, such as transposases, recombinases, integrases, or phage proteins (Figure 1E). Of these, 79 homologs had a sequence similarity of >99% with at least one other homolog, 27 homologs had a proportion of G and C bases that significantly differed from the rest of the genome of the isolate (Figure 1F). Finally, 8 homologs were very strong candidates for HGT, displaying a >99% sequence similarity with at least one other homolog and having a GC content that significantly differed from the rest of the genome of the isolate, representing 0.3% of all RND-antiporter homologs of this study.

RND-antiporter homologs of leaf bacteria clustered together in the phylogenetic tree more often than their counterparts from root bacteria, resulting in 14 clusters of > 30 homologs from leaf-associated isolates (thereafter called “LC” for “leaf cluster”), versus none in root-associated isolates (Figure 1B). We looked at the HGT markers within these clusters, speculating that some clusters might be the result of HGT (Figure 1F). Strikingly, all 8 homologs identified above with strong HGT markers (flanked with genome plasticity genes, similarity of >99% and different GC content from the rest of the genome) were found in LC9, and over half (55%) of the homologues in this cluster exhibited at least one HGT marker, suggesting an evolutionary advantage in gaining an antiporter homolog from this heavy-metal-exporting cluster (Figure 1F). The other clusters also exhibited some HGT markers, although less prevalent (Figure 1F). Overall, this points to a lesser diversification of homologs from leaf isolates in some subclades, and to an increase in horizontally transferred genes in some heavy-metal exporting subclades.

### Sequence and structure conservation of RND-antiporters differs by domain

One very interesting feature of RND-antiporters is their relatively low sequence conservation. Overall, the amino acid sequence conservation of RND-antiporters was quite low, with an average 41% sequence conservation across all residues (Figure 2A). This diversification was elevated between members of different clades (with pairwise similarity sometimes as low as 10.7%), suggesting either strong diversifying selection over the entire sequence or a long evolutionary time since the divergence of the clades (Supplementary Figure S3). This high degree of diversification is in stark contrast to the largely conserved structure of RND antiporters, which one might expect to be a product of sequence conservation. Considering the high level of diversity and the need for functional conservation, we hypothesized that the protein may contain functionally essential and consequently more conserved regions, forming a structural scaffold, and regions that are subjected to relaxed or even diversifying selection and consequently more diversified located in the specificity conferring regions of the protein.

In order to distinguish between more conserved and more diversified regions of the proteins, we investigated to what extend different protein sub-domains (Kobylka et al., 2020; Murakami et al., 2002) exhibited different degrees of conservation (Figure 2A, B, see M&M Consensus sequences, structure prediction and analysis for details). We used the consensus sequence of each clade to predict consensus protein structures with AlphaFold3 (Abramson et al., 2024), and assigned each residue to a protein sub-domain based on the comparison of the consensus structure with the crystal structure of AcrB (Murakami et al., 2002).

We then calculated sequence conservation for each of the different domains of RND-antiporters (Figure 2A, Supplementary Figure S3, see M&M Consensus sequence, structure prediction and analysis for details). Sequence conservation varied depending on the protein domain, with transmembrane domains being the most conserved (Figure 2A). Notably, the transmembrane domain 7 was less conserved than other transmembrane domains, with a per-clade conservation of 37-48%, while the transmembrane domain 4 showed the highest conservation, with a per-clade conservation of 62-83% (Figure 2A). The porter and funnel domains were less conserved. This may be explained by the structural restrictions imposed on transmembrane domains (Oberai et al., 2009) leading to a higher level of conservation compared to the porter and funnel domains, which harbor more diversity, and determine the substrate specificity of the antiporter and thus the whole RND transporter (Tam et al., 2021).

As a measure of structural diversity, we calculated the root mean-square deviation (RMSD) of each position’s residue from the predicted structure of each clade’s consensus sequence compared to the predicted structure of the overall consensus (Figure 2C). As for the sequence diversity (Figure 2A), the RMSD values are variable across domains. As expected, the sequence-conserved transmembrane domains are structurally conserved, with low RMSD values, while the porter and funnel domains are more structurally diverse. Different clades showed different RMSD patterns, with Clades I, II, III, and V displaying structural divergence of PC2 domains (4.8-7.1 Å) and Clade VI showing an RMSD peak of 15Å in the PC1 domain (Figure 2C). All clades had RMSD values above 5Å at the interface between transmembrane domains 6 and 7, corresponding to the Iɑ helix facing the cytoplasm. Altogether, this confirms that transmembrane domains are structurally conserved across all clades, and that each clade shows specific structural features, differing between clades and from the general consensus structure. These features are mainly found in the porter and funnel domains, displaying a higher structural (Figure 2C) and sequence (Figure 2A) diversification. Taken together this suggests variation of the substrate recognition and transport domains enabling consequential changes in substrate specificity possibly reflecting adaptations to the plant environment.

We calculated the clade-specific nonsynonymous substitution rate (β) and the synonymous substitution rate (ɑ) for each residue of all RND-antiporter homologs (Figure 2D, Supplementary Figure S4). We classify the type of diversification using the β-ɑ metric, which is more reliable than the classic dN/dS (ie. β/ɑ), as this estimate is prone to errors when ɑ is small (Murrell et al., 2013). There was no signature of diversifying selection (β-ɑ > 0) in clades I, III and V, and two instances of mild diversifying selection in clades II and IV, one single residue in each clade (Figure 2D, E). Due to high sequence diversity, we could not calculate substitution rates for Clade VI and therefore excluded it from the rest of the analysis. Most purifying selection (β-ɑ < 0) is relatively mild (around 2.5), but some sites are under stronger purifying selection. Mapping of the diversification metric on predicted protein structures did not show any hotspots of diversifying selection on a particular region (Figure 2E), suggesting that the amino acids that drive substrate diversity are evolving at the same rate as random mutations, indicating an unconstrained evolution of substrate specificity rather than driven by directional selection.

### RND transporters harbor clade-specific conserved motifs

We looked for motifs that were present in >75% of all homologs of a clade and not present in any homolog of the other clades using the MAFFT package (see M&M Motif discovery, statistical analysis and data visualization). While Clades V and VI do not have clade-specific motifs, all other clades have several clade-specific conserved motifs in the transmembrane domains (Figure 3). The PN1 domains and the PN2-DN interface also show conserved motifs in several clades; Clade I is the only clade with conserved motifs in the transmembrane domain 1 and PC2 domains. This shows that the transmembrane domains are conserved within clades, and that additional clade-specific motifs can be found across the protein structure. Interestingly, very few conserved motifs are found in the putative substrate entry channel (Figure 3, Supplementary Figure S5), strengthening our claim that structural and sequence diversity in this location is important for substrate diversity of RND transporters.

## Discussion

### RND pumps are enriched in plant-associated bacteria

Our initial assessment of RND efflux pumps in plant-associated bacteria confirmed the trend previously observed by others: the plant environment appears to be preferentially colonized by bacteria with a large number of RND efflux pumps. Given the chemical complexity of plant specialized metabolites, adaptation to this ecological niche likely depends on the evolution of multiple detoxification strategies. Adaptation to a specific host may take the form of chemical modifications to host-specific compounds, such as previously described in Sphingomonas species colonizing benzoxazinoid-producing maize (Thoenen et al., 2023). Contrastingly, detoxification via broad-spectrum efflux pumps may thus be a signature of broader adaptation to the plant environment, rather than a more committed adaptation to a specific host lineage (Martinez et al., 2009; Piddock, 2006). Nevertheless, a specific broad-spectrum efflux transporter may still be essential for the establishment in a specific host environment, as exemplified by the ef90 antiporter, previously established to facilitate adaptation to the *A. thaliana* leaf environment. The presence of close homologs of the ef90 antiporter in our dataset supports the role of this particular antiporter in Arabidopsis leaf colonization, particularly given that the strains harboring these homologs have been isolates from soils collected at sites over 7.000km away from each other (Bai et al., 2015; Finkel et al., 2020).

Taken together, enrichment of broad-spectrum RND efflux transporters may promote general adaptation to host associated niches, thereby contributing to the previously described core plant microbiota (Hacquard et al., 2015).

### RND antiporters form monophyletic clades reflecting substrate specificity

In our analysis, we found 231 plant-associated isolates harboring 7 or more homologs per genome, representing 80% of the 282 plant associated isolates analyzed (Supplementary Table S1), indicating that plant-associated bacteria genomes harbor a large diversity of RND substrate specificities, and more importantly, expansions in multiple phylogenetic clades (Supplementary Table S2). Individual genomes harbored between 1 and 26 RND-antiporter homolog(s), with an overall average of 10.5 homologs per genome (Figure 1A). This can be rounded to the number reported earlier for Arabidopsis root-associated Gram-negative bacteria harbored on average 11 RND-antiporter copies (Russ et al., 2025). Despite the overall high copy number of RND antiporters identified in plant-associated isolates, there are substantial differences in the distribution of antiporters across different bacterial taxa (Supplementary Table S3). This finding suggests that the diversification of the RND antiporters likely precedes the diversification of Proteobacteria, and these efflux systems subsequently evolved within different bacterial lineages while retaining core genetic advantages for host adaptation. We identified six monophyletic clades of antiporter homologs, with a clear separation of the previously described HAE-1, NFE and HME clades. While the clades did not separate by organ of origin, homologs from leaf-associated bacteria tended to cluster together, indicating a higher degree of conservation or selection in specific environmental niches.

### Horizontal transfer of RND components

The HAE-1 clade, which contained the majority (63.9%) of RND antiporter homologs, displayed indicators of being involved in adaptation to the leaf environment. However, an exploration of potential signatures of recent horizontal transfer did not produce strong signals in this clade. Instead, we identified 8 homologs in the HME (heavy metal exporting) clade with support for recent HGT based on the following markers: high DNA sequence similarity between the homologs, GC content of the genes deviating from genomic background, and proximity to flanking genes that are typically associated with mobility or plasticity. While these eight genes displayed strong indications of recent transfer, an investigation of the phylogenetic leaf cluster (LC9) which contains these genes revealed that 55% of the genes in this clade were positive for at least one of the HGT markers. LC9 thus seems to contain RND components with increased genomic mobility, correlated with the presence of members of this clade in bacteria isolated from leaves. Given that we do not know the function of the 8 homologs with strong indications of being recently transferred, we cannot unambiguously attribute their spread across genera in these populations to a selective advantage or to the propagation of a selfish genetic element, spreading independently of host fitness benefits.

### Expansion of the NFE clade

The large expansion of the NFE clade observed in *A. thaliana*-associated bacteria was driven by homologs from multiple bacterial orders not typically associated with root nodule symbiosis (Table 1, Supplementary Figure S1A, Table S4). The NFE clade is assumed to export Lipo-chitooligosaccharides (LCO), which are important for the initiation of root nodule symbiosis (Dénarié et al., 1996). LCO production has been detected not only in rhizobial bacteria but also across widely divergent fungi lineages, including taxa not known to form plant symbioses, and has been proposed to regulate fungi growth and development (Rush et al., 2020). Moreover, LCO-analogous activity has been reported in the non-rhizobia bacterium *Bacillus circulans* (Lian et al., 2001). Therefore, the putative function of RND pumps in the NFE clade for LCO transport may either be a signal that LCO production and transport are much more widespread than previously assumed (Dénarié et al., 1996), or this clade transports related substrates with broad distribution and yet unknown function potentially related to organismic interactions.

### Structural conservation in the absence of sequence conservation

The antiporter component of the RND complex displays remarkably low sequence conservation, at both nucleotide and amino acid sequence levels. Considering the physicochemical interactions of the antiporter component with the adaptor linking to the channel, some degree of structural conservation is likely required to maintain these interactions (Daury et al., 2016). At the same time, structural diversification of the sites responsible for substrate specificity may be required to avoid reaching an adaptive dead-end, where hyper-specialization to a specific chemical environment may not be desirable with the perspective of fluctuating chemical environments encountered during the shift between plant hosts. Even on a single host, the niche-specificity can give rise to frequent metabolite variations, especially in exposed leaves responding to a variety of biotic and abiotic stresses (Sasse et al., 2018; Vorholt, 2012). Thus, it is conceivable that the substrate promiscuity observed in RND transporters is an intrinsic feature of this class. This line of thought is further supported by a general failure to unambiguously identify substrate binding and entry sites in the antiporter component (Du et al., 2018; Wilhelm and Pos, 2024), potentially indicating flexibility in substrate uptake. Taking this further, one might speculate that the specificity of RND transporters is perhaps maintained negatively, with selection working towards preventing the leakage of important resources, rather than selecting for export of specific substrates. This would distribute specificities across the larger population, allowing for standing diversity in substrate specificities across a population, in turn enabling fast adaptation in the case of a specific, novel chemical stress. Nevertheless, structure gives rise to function, and mapping which sites, or structural elements, drive substrate specificity is important future work.

## Acknowledgements

Molecular graphics and analyses performed with UCSF ChimeraX, developed by the Resource for Biocomputing, Visualization, and Informatics at the University of California, San Francisco, with support from National Institutes of Health R01-GM129325 and the Office of Cyber Infrastructure and Computational Biology, National Institute of Allergy and Infectious Diseases. Thank you to Sergei Pond and colleagues for providing GitHub repositories with FUBAR scripts. This project was funded by the Deutsche Forschungsgemeinschaft (DFG) – in the context of SPP 2125: Dekonstruktion und Rekonstruktion der pflanzlichen Mikrobiota “DECRyPT” project numbers 401867691 and 466385132 to MP and NS, respectively. High performance computing was performed on the BioHPC hosted at the Leibniz Rechenzentrum Munich funded by the German Research Foundation (grant INST 86/2050-1 FUGG), regular data analysis was facilitated by funding via the data management project (Inf (I01)) of TRR356 for RStudio compute resources. (DFG project number 491090170). YZ acknowledges the support by the China Scholarship Council program. LR and YZ are grateful to the Life Science Munich Graduate program for their support.

## Author contributions

### Project management

The project was conceived by all authors. MP identified AcrB homologs as the transporter class of interest for this study. NS and MP acquired funding for the conducted work and provisioned necessary resources, supervised the work and guided experimental design and interpretation.

### Manuscript writing

All authors drafted and revised the manuscript.

### Data generation and analysis

LR curated genomes (Tab S1), extracted homologs, created alignments, constructed phylogenetic trees, implemented and conducted the analysis of markers for horizontal transfer (Figure 1, S2), performed structural predictions and analysis thereof, as well as inference of selection (Figures 2, S3, S4), and carried out the analysis of conserved motifs (Figures 3, S5). YZ contributed to data validation of the AcrB homology search, curated previous results and carried out the comparisons to previous results (Tab 1, S2, S3, S4, S5) and carried out data-visualization for Figure S1A. The table below summarizes which author was responsible for which aspect of the analysis.

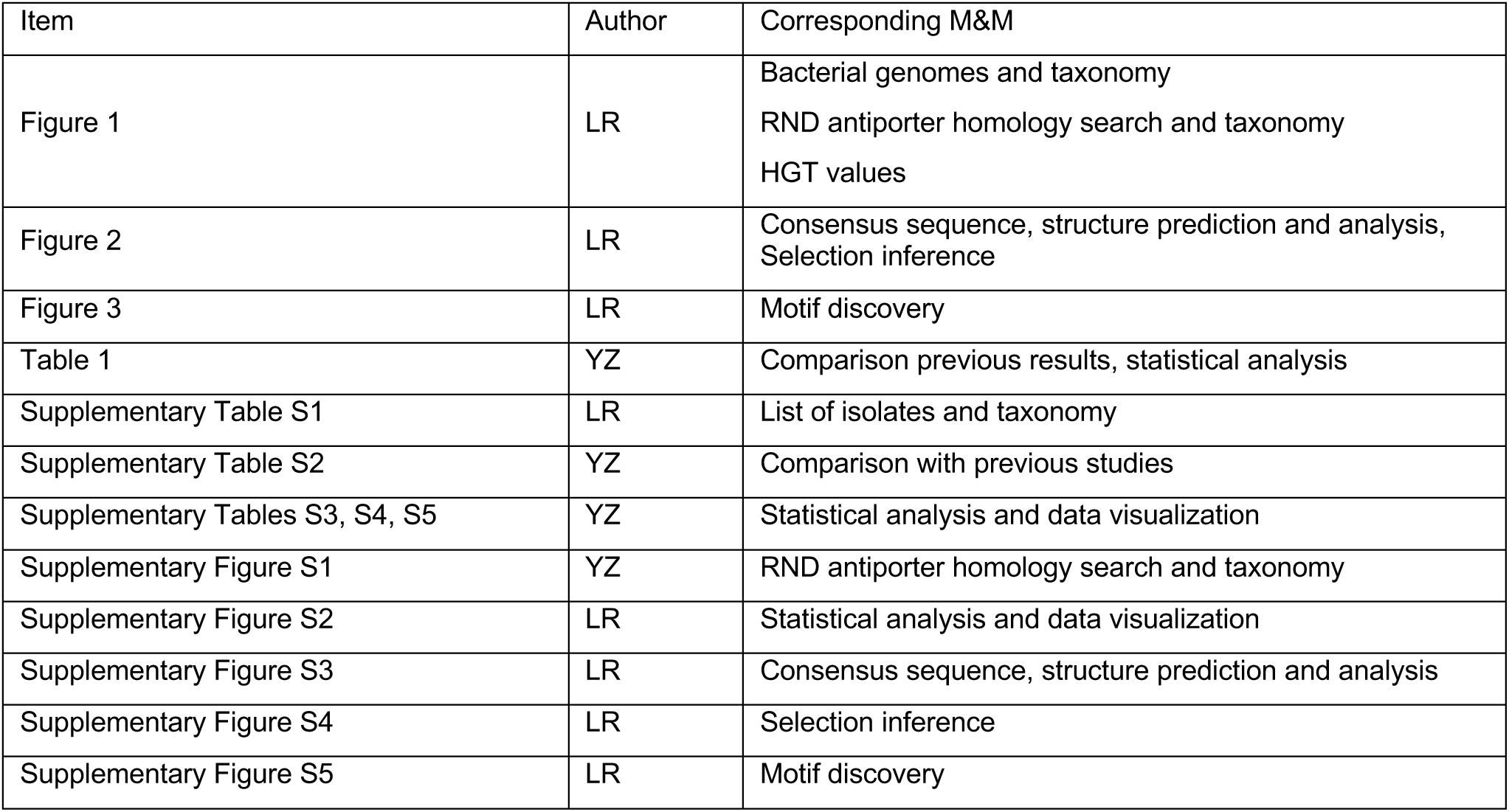

| Item | Author | Corresponding M&M |
| --- | --- | --- |
| Figure 1 | LR | Bacterial genomes and taxonomy<br>RND antiporter homology search and taxonomy<br>HGT values |
| Figure 2 | LR | Consensus sequence, structure prediction and analysis,<br>Selection inference |
| Figure 3 | LR | Motif discovery |
| Table 1 | YZ | Comparison previous results, statistical analysis |
| Supplementary Table S1 | LR | List of isolates and taxonomy |
| Supplementary Table S2 | YZ | Comparison with previous studies |
| Supplementary Tables S3, S4, S5 | YZ | Statistical analysis and data visualization |
| Supplementary Figure S1 | YZ | RND antiporter homology search and taxonomy |
| Supplementary Figure S2 | LR | Statistical analysis and data visualization |
| Supplementary Figure S3 | LR | Consensus sequence, structure prediction and analysis |
| Supplementary Figure S4 | LR | Selection inference |
| Supplementary Figure S5 | LR | Motif discovery |

## Supplementary material

**Supplementary Figure S1.**
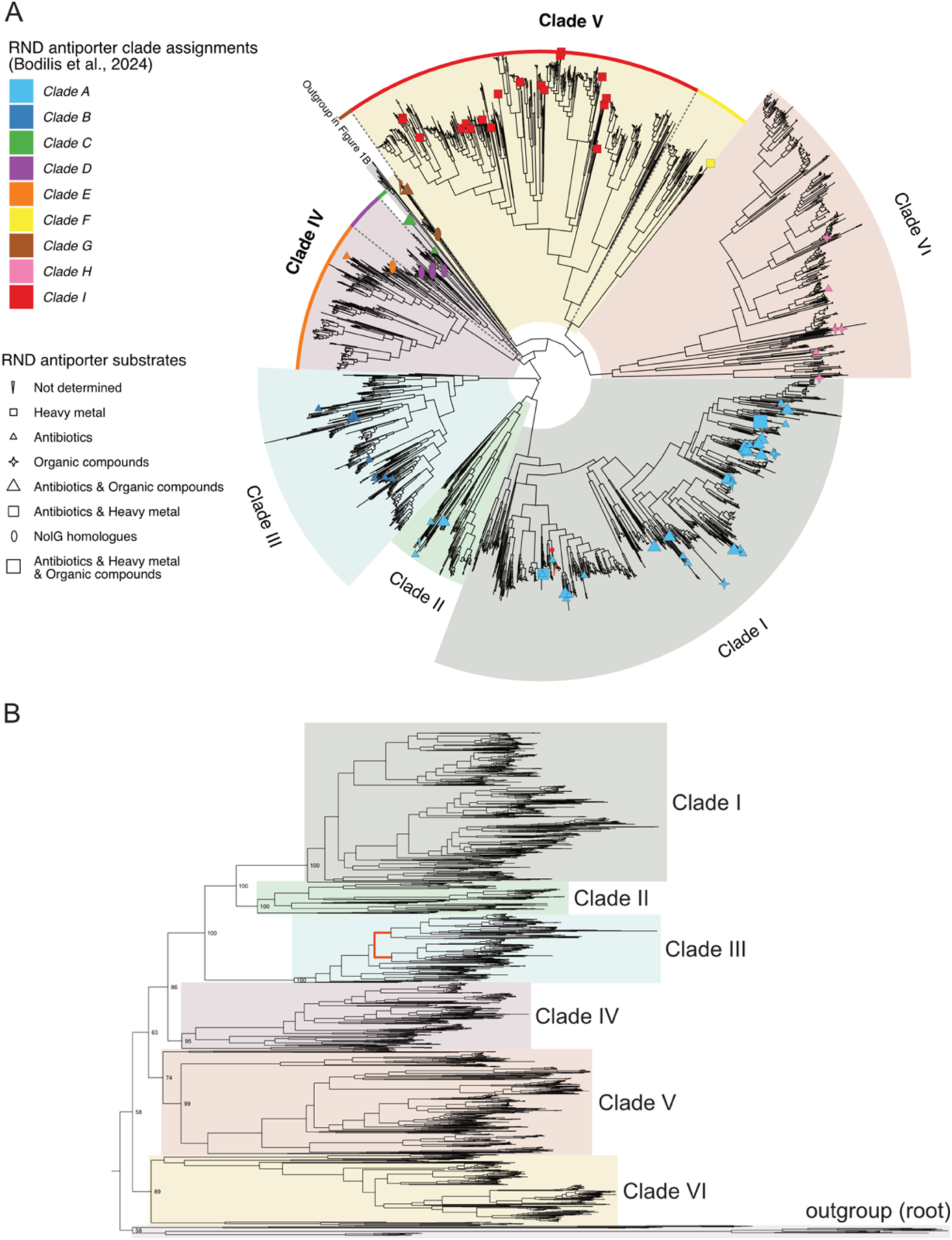
Phylogenetic classification of the RND antiporter homologs in this study and comparison with previous reports. **A.** Translating the clade nomenclature of independent RND antiporter phylogenetic studies using functionally defined RND antiporters as Rosetta stone. Phylogenetic tree of RND antiporters identified in this study (AtSphere) and from Transporter Classification Database (TCDB) with characterized functions. TCDB antiporters are distinguished by unique shapes corresponding to their known substrate specificities. The color of each shape represents clade assignments based on the classification by Bodilis *et al*. (2024). Notably, Clades IV and VI (bolded) in this study comprise multiple monophyletic groups previously defined by Bodilis *et al*. (2024), which are highlighted with colored arcs to reflect their previous classification. The 21 homologs showing >84% sequence similarity to the ef90 antiporter reported by Russ et al. (2025) are highlighted as green (leaf isolates) and red (root isolates) branches, with a single red node marking the subcluster. **B.** Phylogenetic tree of RND-antiporters from AtSphere with bootstrap values for each clade. The single ancestral duplication reported by Bodilis *et al*. 2024 is highlighted in red. This tree is additional information for Figure 1B.

**Supplementary Figure S2:**
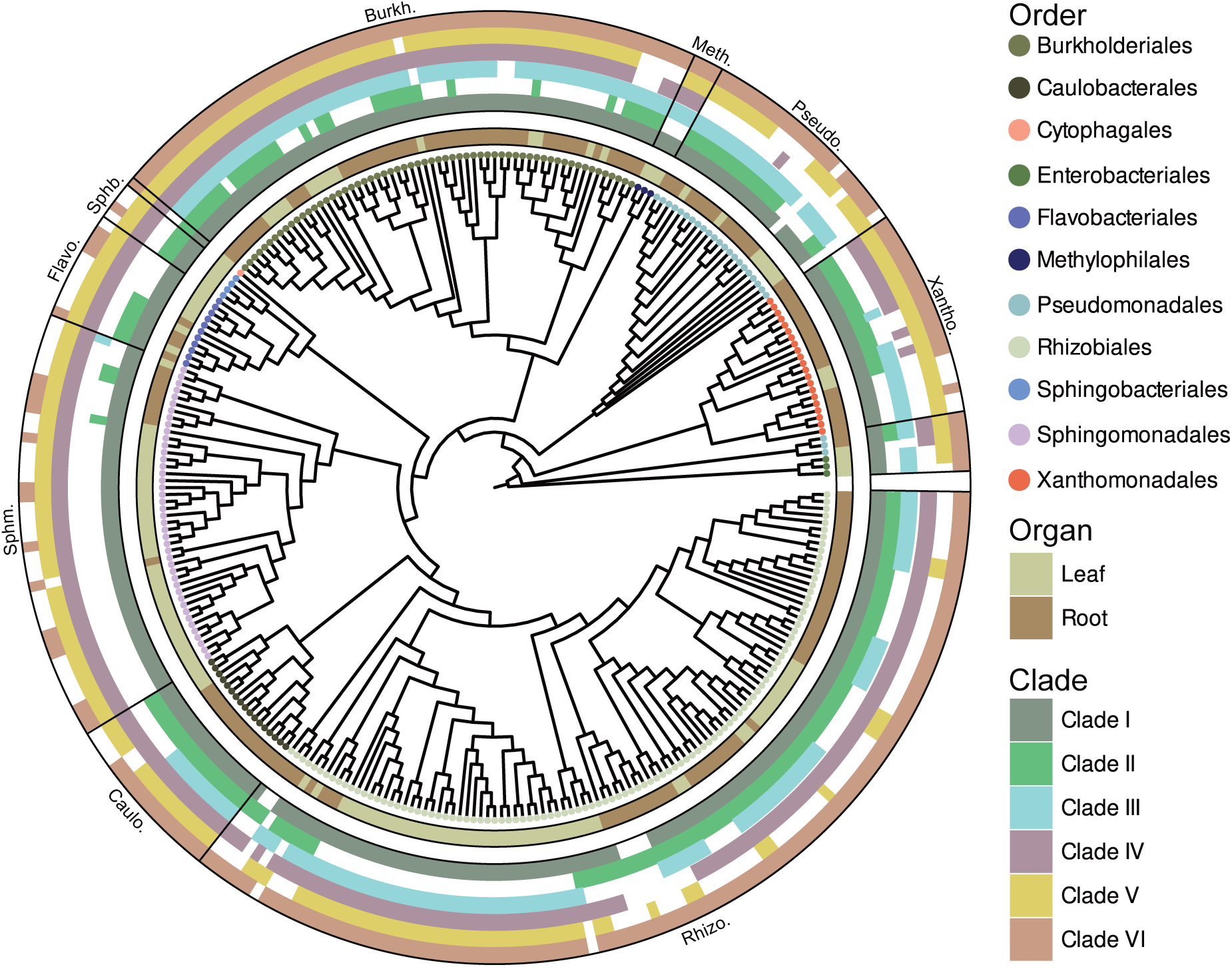
Phylogenetic tree of the isolates used in this study showing the presence/absence of homologs from each clade for each isolate.

**Supplementary Figure 3.**
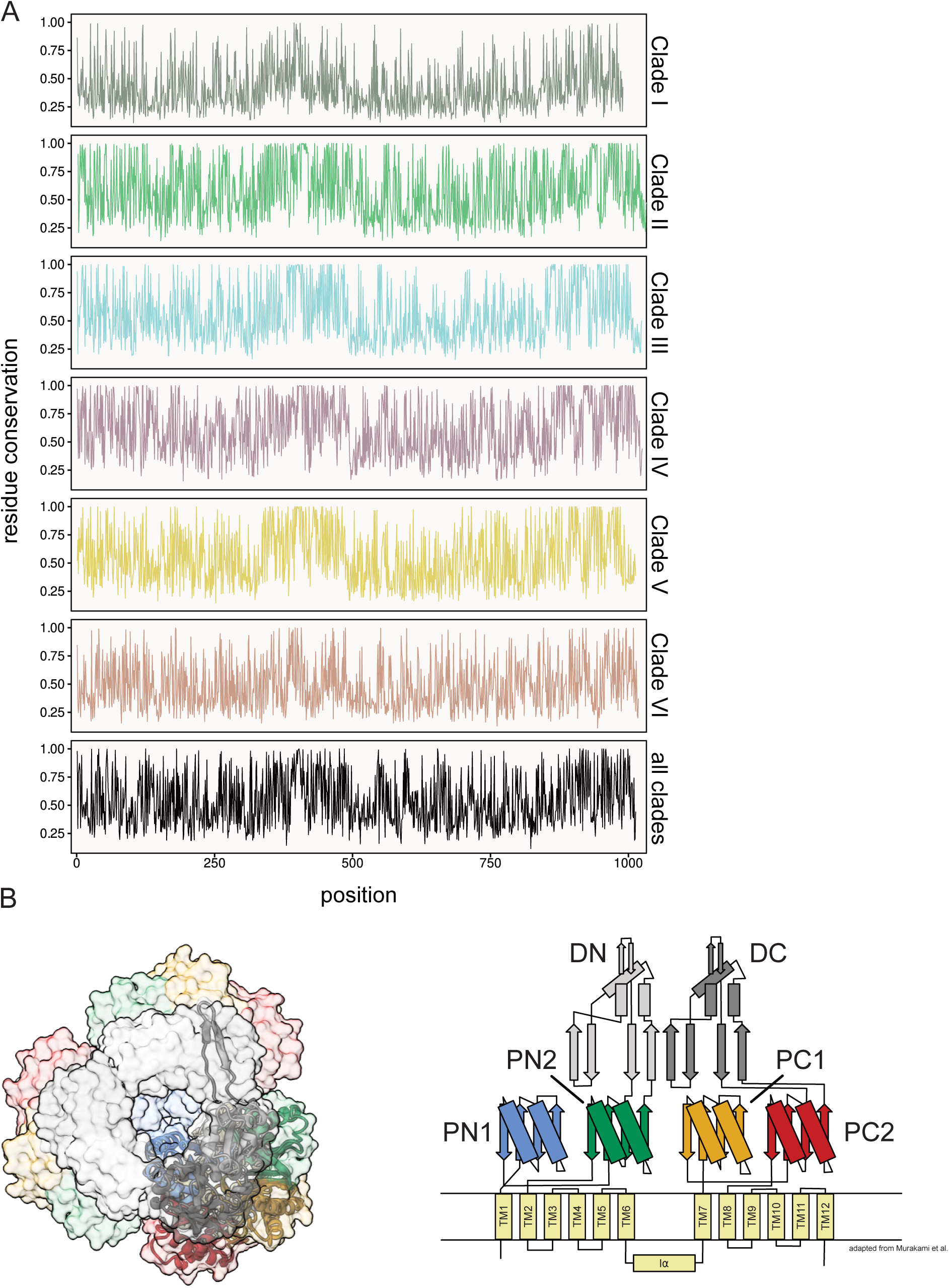
RND antiporter residue conservation and domains. **A.** Conservation of each residue of RND antiporters across each clade and across all homologs (bottom). **B.** Predicted trimeric protein structure of the consensus sequence of all RND permeases homologues, colored by domains, viewed from the top (top) and schematic representation of the RND permease domains (bottom; adapted from Murakami et al., 2002).

**Supplementary Figure S4.**
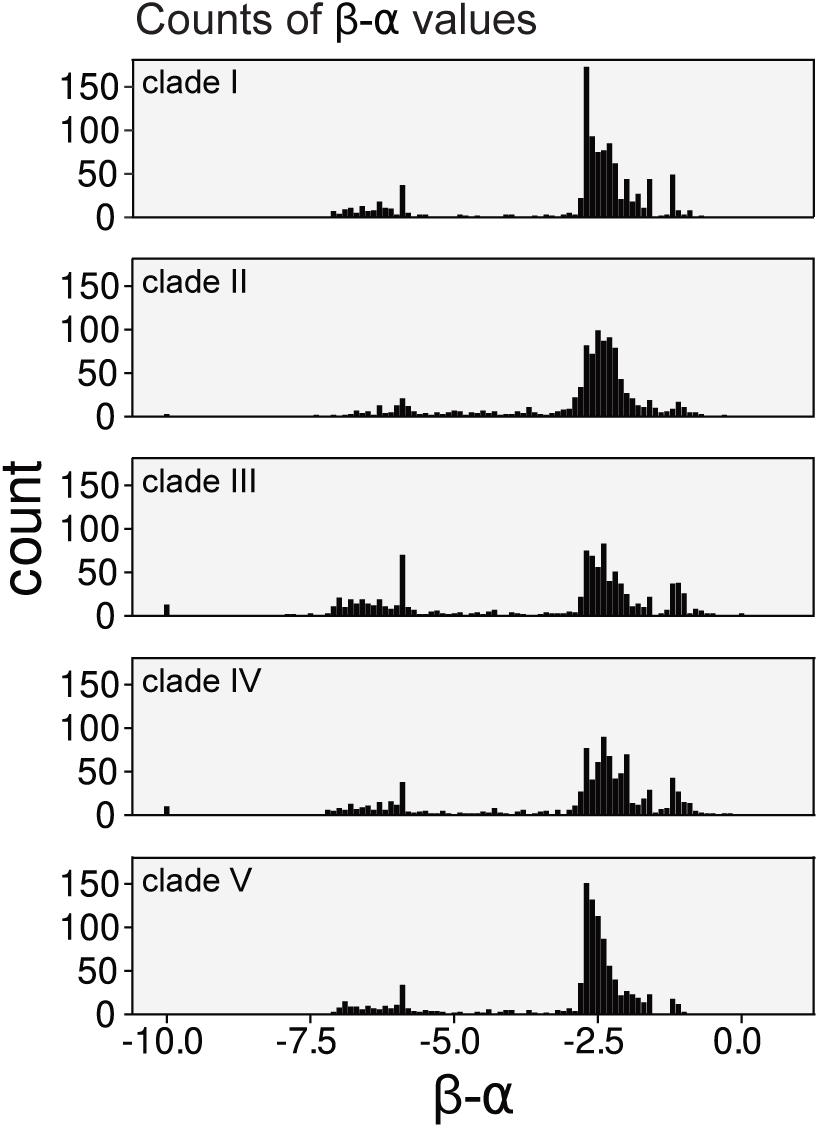
Distribution of selection values in each clade. Distribution of β-α values across all residues of clades A-E, with an artificial minimum at -10, calculated for each individual residue. Negative β-α indicates purifying selection. Protein structure predictions were done with AlphaFold3 on the AlphaFold Server.

**Supplementary Figure S5.**
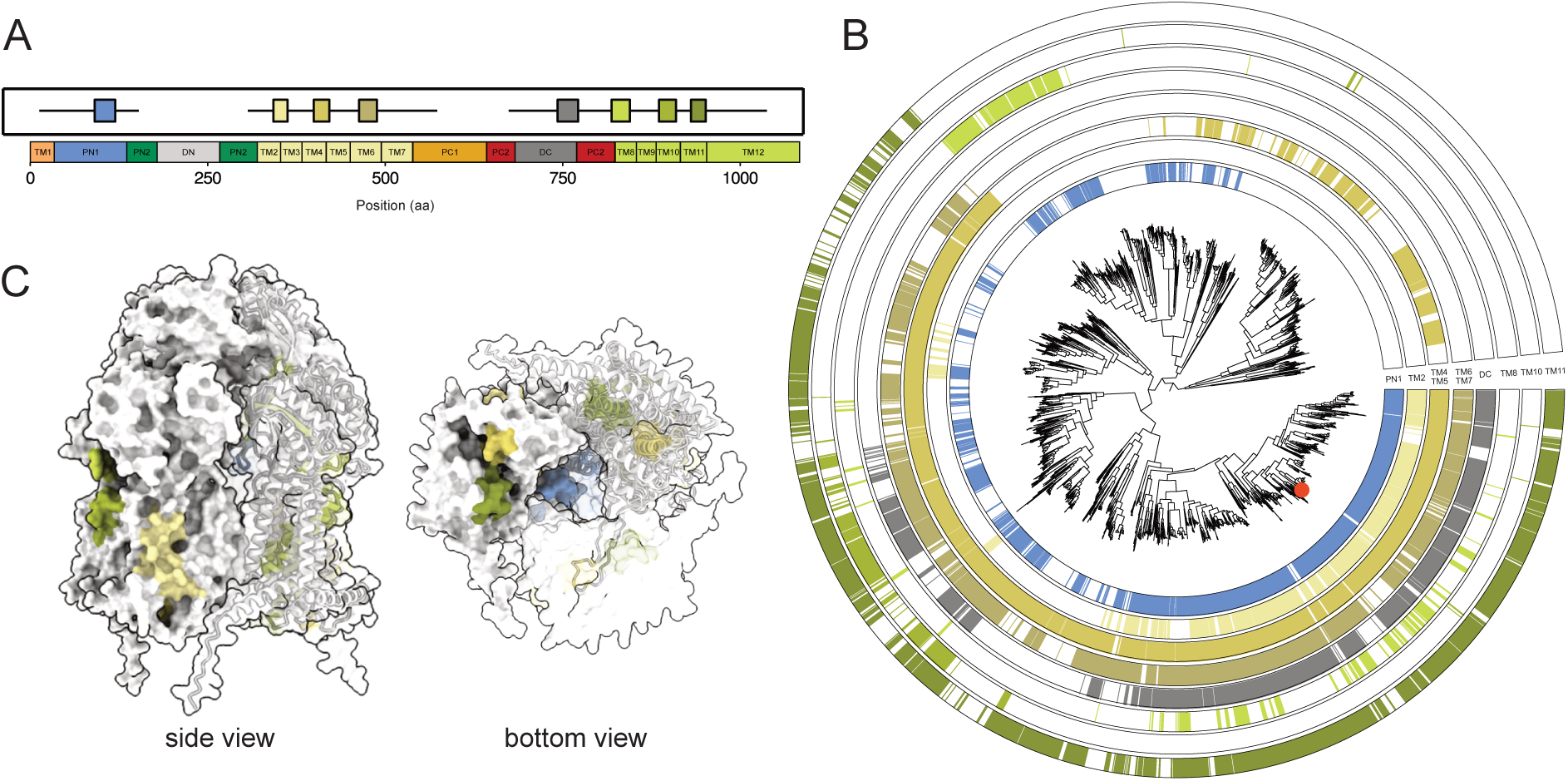
Conservation of motifs across all clades. **A.** Localization of motifs conserved across RND transporter homologs from all clades. **B.** Distribution of conserved motifs. **C.** Conserved motif coloured on the predicted protein structure of one RND antiporter homolog harbouring all conserved motifs, side view (left) and bottom view (right). The homolog is indicated with a red dot on the phylogenetic tree in B.

**Supplementary Table S1.** Taxonomy of isolates used for RND antiporter homologue screening count of RND antiporters per isolate.

| Isolate | Organ | Order | Homologue count |
| --- | --- | --- | --- |
| Root436 | Root | Rhizobiales | 1 |
| Root635 | Root | Rhizobiales | 2 |
| Root105 | Root | Rhizobiales | 3 |
| Root1471 | Root | Rhizobiales | 3 |
| Root413D1 | Root | Rhizobiales | 3 |
| Root554 | Root | Rhizobiales | 3 |
| Root569 | Root | Pseudomonadales | 3 |
| Leaf168 | Leaf | Caulobacterales | 4 |
| Leaf280 | Leaf | Caulobacterales | 4 |
| Leaf344 | Leaf | Rhizobiales | 4 |
| Leaf363 | Leaf | Caulobacterales | 4 |
| Leaf412 | Leaf | Sphingomonadales | 4 |
| Root1279 | Root | Caulobacterales | 4 |
| Root608 | Root | Caulobacterales | 4 |
| Leaf10 | Leaf | Sphingomonadales | 5 |
| Leaf17 | Leaf | Sphingomonadales | 5 |
| Leaf180 | Leaf | Flavobacteriales | 5 |
| Leaf20 | Leaf | Sphingomonadales | 5 |
| Leaf22 | Leaf | Sphingomonadales | 5 |
| Leaf231 | Leaf | Sphingomonadales | 5 |
| Leaf24 | Leaf | Sphingomonadales | 5 |
| Leaf25 | Leaf | Sphingomonadales | 5 |
| Leaf33 | Leaf | Sphingomonadales | 5 |
| Leaf34 | Leaf | Sphingomonadales | 5 |
| Leaf407 | Leaf | Sphingomonadales | 5 |
| Leaf42 | Leaf | Sphingomonadales | 5 |
| Leaf420 | Leaf | Rhizobiales | 5 |
| Leaf5 | Leaf | Sphingomonadales | 5 |
| Leaf64 | Leaf | Rhizobiales | 5 |
| Leaf67 | Leaf | Sphingomonadales | 5 |
| Root157 | Root | Rhizobiales | 5 |
| Root672 | Root | Sphingomonadales | 5 |
| Root685 | Root | Rhizobiales | 5 |
| Root710 | Root | Sphingomonadales | 5 |
| Leaf11 | Leaf | Sphingomonadales | 6 |
| Leaf16 | Leaf | Sphingomonadales | 6 |
| Leaf226 | Leaf | Sphingomonadales | 6 |
| Leaf23 | Leaf | Sphingomonadales | 6 |
| Leaf29 | Leaf | Sphingomonadales | 6 |
| Leaf30 | Leaf | Sphingomonadales | 6 |
| Leaf32 | Leaf | Sphingomonadales | 6 |
| Leaf343 | Leaf | Sphingomonadales | 6 |
| Leaf41 | Leaf | Sphingobacteriales | 6 |
| Leaf443 | Leaf | Rhizobiales | 6 |
| Leaf62 | Leaf | Sphingomonadales | 6 |
| Leaf9 | Leaf | Sphingomonadales | 6 |
| Root1497 | Root | Sphingomonadales | 6 |
| Root154 | Root | Sphingomonadales | 6 |
| Root214 | Root | Sphingomonadales | 6 |
| Root241 | Root | Sphingomonadales | 6 |
| Root552 | Root | Rhizobiales | 6 |
| Leaf194 | Leaf | Sphingobacteriales | 7 |
| Leaf208 | Leaf | Sphingomonadales | 7 |
| Leaf324 | Leaf | Rhizobiales | 7 |
| Leaf341 | Leaf | Rhizobiales | 7 |
| Leaf371 | Leaf | Rhizobiales | 7 |
| Leaf38 | Leaf | Sphingomonadales | 7 |
| Leaf384 | Leaf | Rhizobiales | 7 |
| Leaf394 | Leaf | Flavobacteriales | 7 |
| Root100 | Root | Rhizobiales | 7 |
| Root1294 | Root | Sphingomonadales | 7 |
| Root1423 | Root | Caulobacterales | 7 |
| Root482 | Root | Rhizobiales | 7 |
| Root50 | Root | Sphingomonadales | 7 |
| Root720 | Root | Sphingomonadales | 7 |
| Leaf129 | Leaf | Pseudomonadales | 8 |
| Leaf155 | Leaf | Rhizobiales | 8 |
| Leaf176 | Leaf | Sphingobacteriales | 8 |
| Leaf201 | Leaf | Flavobacteriales | 8 |
| Leaf242 | Leaf | Sphingomonadales | 8 |
| Leaf262 | Leaf | Rhizobiales | 8 |
| Leaf28 | Leaf | Sphingomonadales | 8 |
| Leaf357 | Leaf | Sphingomonadales | 8 |
| Leaf383 | Leaf | Rhizobiales | 8 |
| Leaf400 | Leaf | Burkholderiales | 8 |
| Leaf404 | Leaf | Flavobacteriales | 8 |
| Leaf405 | Leaf | Flavobacteriales | 8 |
| Leaf454 | Leaf | Rhizobiales | 8 |
| MF033 | Root | Rhizobiales | 8 |
| Root1212 | Root | Rhizobiales | 8 |
| Root1240 | Root | Rhizobiales | 8 |
| Root209 | Root | Burkholderiales | 8 |
| Root268 | Root | Rhizobiales | 8 |
| Root274 | Root | Rhizobiales | 8 |
| Root494 | Root | Xanthomonadales | 8 |
| Root901 | Root | Flavobacteriales | 8 |
| CL041 | Root | Rhizobiales | 9 |
| Leaf198 | Leaf | Sphingomonadales | 9 |
| Leaf2 | Leaf | Sphingomonadales | 9 |
| Leaf202 | Leaf | Rhizobiales | 9 |
| Leaf216 | Leaf | Sphingobacteriales | 9 |
| Leaf230 | Leaf | Sphingomonadales | 9 |
| Leaf311 | Leaf | Rhizobiales | 9 |
| Leaf321 | Leaf | Rhizobiales | 9 |
| Leaf359 | Leaf | Flavobacteriales | 9 |
| Leaf386 | Leaf | Rhizobiales | 9 |
| Leaf391 | Leaf | Rhizobiales | 9 |
| Leaf427 | Leaf | Rhizobiales | 9 |
| Leaf453 | Leaf | Rhizobiales | 9 |
| Leaf460 | Leaf | Rhizobiales | 9 |
| Leaf68 | Leaf | Rhizobiales | 9 |
| Leaf78 | Leaf | Burkholderiales | 9 |
| MF224 | Root | Rhizobiales | 9 |
| Root1204 | Root | Rhizobiales | 9 |
| Root1277 | Root | Caulobacterales | 9 |
| Root1290 | Root | Caulobacterales | 9 |
| Root381 | Root | Rhizobiales | 9 |
| Root420 | Root | Flavobacteriales | 9 |
| Root480 | Root | Xanthomonadales | 9 |
| Root483D1 | Root | Rhizobiales | 9 |
| Root491 | Root | Rhizobiales | 9 |
| Root65 | Root | Xanthomonadales | 9 |
| Root670 | Root | Rhizobiales | 9 |
| Root77 | Root | Caulobacterales | 9 |
| Leaf127 | Leaf | Pseudomonadales | 10 |
| Leaf160 | Leaf | Burkholderiales | 10 |
| Leaf26 | Leaf | Sphingomonadales | 10 |
| Leaf306 | Leaf | Rhizobiales | 10 |
| Leaf434 | Leaf | Pseudomonadales | 10 |
| Leaf50 | Leaf | Enterobacteriales | 10 |
| Leaf51 | Leaf | Enterobacteriales | 10 |
| Leaf53 | Leaf | Enterobacteriales | 10 |
| Leaf82 | Leaf | Flavobacteriales | 10 |
| Root102 | Root | Rhizobiales | 10 |
| Root1334 | Root | Rhizobiales | 10 |
| Root142 | Root | Rhizobiales | 10 |
| Root1455 | Root | Caulobacterales | 10 |
| Root1462 | Root | Rhizobiales | 10 |
| Root172 | Root | Rhizobiales | 10 |
| Root267 | Root | Burkholderiales | 10 |
| Root275 | Root | Burkholderiales | 10 |
| Root329 | Root | Pseudomonadales | 10 |
| Root402 | Root | Burkholderiales | 10 |
| Root405 | Root | Burkholderiales | 10 |
| Root483D2 | Root | Rhizobiales | 10 |
| Root655 | Root | Caulobacterales | 10 |
| Root662 | Root | Burkholderiales | 10 |
| Root695 | Root | Rhizobiales | 10 |
| Root700 | Root | Caulobacterales | 10 |
| Root73 | Root | Rhizobiales | 10 |
| Soil773 | Root | Xanthomonadales | 10 |
| Leaf130 | Leaf | Pseudomonadales | 11 |
| Leaf148 | Leaf | Xanthomonadales | 11 |
| Leaf191 | Leaf | Burkholderiales | 11 |
| Leaf220 | Leaf | Burkholderiales | 11 |
| Leaf274 | Leaf | Burkholderiales | 11 |
| Leaf48 | Leaf | Pseudomonadales | 11 |
| Leaf76 | Leaf | Burkholderiales | 11 |
| Leaf84 | Leaf | Burkholderiales | 11 |
| Root1280 | Root | Pseudomonadales | 11 |
| Root179 | Root | Xanthomonadales | 11 |
| Root186 | Root | Flavobacteriales | 11 |
| Root189 | Root | Burkholderiales | 11 |
| Root217 | Root | Burkholderiales | 11 |
| Root318D1 | Root | Burkholderiales | 11 |
| Root423 | Root | Rhizobiales | 11 |
| Root473 | Root | Burkholderiales | 11 |
| Root487D2Y | Root | Caulobacterales | 11 |
| Root558 | Root | Rhizobiales | 11 |
| Root561 | Root | Xanthomonadales | 11 |
| Root562 | Root | Pseudomonadales | 11 |
| Root564 | Root | Rhizobiales | 11 |
| Root604 | Root | Xanthomonadales | 11 |
| Root656 | Root | Caulobacterales | 11 |
| Root68 | Root | Pseudomonadales | 11 |
| Root70 | Root | Burkholderiales | 11 |
| Root71 | Root | Pseudomonadales | 11 |
| Root916 | Root | Xanthomonadales | 11 |
| Root935 | Root | Flavobacteriales | 11 |
| Root983 | Root | Xanthomonadales | 11 |
| Leaf139 | Leaf | Burkholderiales | 12 |
| Leaf15 | Leaf | Pseudomonadales | 12 |
| Leaf267 | Leaf | Burkholderiales | 12 |
| Leaf469 | Leaf | Rhizobiales | 12 |
| Leaf58 | Leaf | Pseudomonadales | 12 |
| MF051 | Root | Pseudomonadales | 12 |
| MF295 | Root | Burkholderiales | 12 |
| MF349 | Root | Burkholderiales | 12 |
| Root1220 | Root | Rhizobiales | 12 |
| Root1252 | Root | Rhizobiales | 12 |
| Root127 | Root | Rhizobiales | 12 |
| Root1298 | Root | Rhizobiales | 12 |
| Root1312 | Root | Rhizobiales | 12 |
| Root1472 | Root | Caulobacterales | 12 |
| Root231 | Root | Rhizobiales | 12 |
| Root258 | Root | Rhizobiales | 12 |
| Root278 | Root | Rhizobiales | 12 |
| Root31 | Root | Rhizobiales | 12 |
| Root335 | Root | Burkholderiales | 12 |
| Root342 | Root | Caulobacterales | 12 |
| Root343 | Root | Caulobacterales | 12 |
| Root401 | Root | Pseudomonadales | 12 |
| Root411 | Root | Burkholderiales | 12 |
| Root434 | Root | Burkholderiales | 12 |
| Root568 | Root | Burkholderiales | 12 |
| Root627 | Root | Xanthomonadales | 12 |
| Root651 | Root | Rhizobiales | 12 |
| Root708 | Root | Rhizobiales | 12 |
| Root74 | Root | Rhizobiales | 12 |
| Root83 | Root | Burkholderiales | 12 |
| Root954 | Root | Rhizobiales | 12 |
| Leaf106 | Leaf | Rhizobiales | 13 |
| Leaf108 | Leaf | Rhizobiales | 13 |
| Leaf131 | Leaf | Xanthomonadales | 13 |
| Leaf265 | Leaf | Burkholderiales | 13 |
| Leaf399 | Leaf | Rhizobiales | 13 |
| Leaf414 | Leaf | Methylophilales | 13 |
| Leaf466 | Leaf | Rhizobiales | 13 |
| Leaf86 | Leaf | Rhizobiales | 13 |
| MF035 | Root | Pseudomonadales | 13 |
| MF048 | Root | Pseudomonadales | 13 |
| MF397 | Root | Pseudomonadales | 13 |
| Root1203 | Root | Rhizobiales | 13 |
| Root1238 | Root | Burkholderiales | 13 |
| Root1272 | Root | Burkholderiales | 13 |
| Root149 | Root | Rhizobiales | 13 |
| Root170 | Root | Burkholderiales | 13 |
| Root219 | Root | Burkholderiales | 13 |
| Root29 | Root | Burkholderiales | 13 |
| Root404 | Root | Burkholderiales | 13 |
| Root559 | Root | Xanthomonadales | 13 |
| Root565 | Root | Burkholderiales | 13 |
| Root667 | Root | Xanthomonadales | 13 |
| Root76 | Root | Xanthomonadales | 13 |
| Root96 | Root | Xanthomonadales | 13 |
| Leaf100 | Leaf | Rhizobiales | 14 |
| Leaf102 | Leaf | Rhizobiales | 14 |
| Leaf112 | Leaf | Rhizobiales | 14 |
| Leaf177 | Leaf | Burkholderiales | 14 |
| Leaf257 | Leaf | Sphingomonadales | 14 |
| Leaf416 | Leaf | Methylophilales | 14 |
| Leaf459 | Leaf | Methylophilales | 14 |
| Leaf85 | Leaf | Rhizobiales | 14 |
| Leaf91 | Leaf | Rhizobiales | 14 |
| Leaf99 | Leaf | Rhizobiales | 14 |
| Root133 | Root | Burkholderiales | 14 |
| Root1480D1 | Root | Burkholderiales | 14 |
| Root16D2 | Root | Burkholderiales | 14 |
| Root351 | Root | Burkholderiales | 14 |
| Root690 | Root | Xanthomonadales | 14 |
| Leaf119 | Leaf | Rhizobiales | 15 |
| Leaf93 | Leaf | Rhizobiales | 15 |
| Leaf94 | Leaf | Rhizobiales | 15 |
| MF036 | Root | Pseudomonadales | 15 |
| Root1217 | Root | Burkholderiales | 15 |
| Root1237 | Root | Burkholderiales | 15 |
| Root123D2 | Root | Rhizobiales | 15 |
| Root1485 | Root | Burkholderiales | 15 |
| Root198D2 | Root | Burkholderiales | 15 |
| Root336D2 | Root | Burkholderiales | 15 |
| Root418 | Root | Burkholderiales | 15 |
| Leaf123 | Leaf | Rhizobiales | 16 |
| Leaf125 | Leaf | Rhizobiales | 16 |
| Leaf189 | Leaf | Cytophagales | 16 |
| Leaf456 | Leaf | Rhizobiales | 16 |
| Leaf465 | Leaf | Rhizobiales | 16 |
| Leaf83 | Leaf | Pseudomonadales | 16 |
| Leaf88 | Leaf | Rhizobiales | 16 |
| Leaf89 | Leaf | Rhizobiales | 16 |
| Leaf92 | Leaf | Rhizobiales | 16 |
| MF045 | Root | Pseudomonadales | 16 |
| Root9 | Root | Pseudomonadales | 16 |
| Leaf104 | Leaf | Rhizobiales | 17 |
| Leaf111 | Leaf | Rhizobiales | 17 |
| Leaf117 | Leaf | Rhizobiales | 17 |
| Leaf126 | Leaf | Burkholderiales | 17 |
| Leaf61 | Leaf | Burkholderiales | 17 |
| Leaf87 | Leaf | Rhizobiales | 17 |
| MF110 | Root | Burkholderiales | 17 |
| Root1444 | Root | Burkholderiales | 17 |
| Leaf113 | Leaf | Rhizobiales | 18 |
| Leaf70 | Leaf | Xanthomonadales | 18 |
| MF278 | Root | Burkholderiales | 18 |
| MF350 | Root | Burkholderiales | 18 |
| Root1221 | Root | Burkholderiales | 18 |
| Root630 | Root | Xanthomonadales | 18 |
| Leaf121 | Leaf | Rhizobiales | 19 |
| Leaf122 | Leaf | Rhizobiales | 19 |
| MF160 | Root | Burkholderiales | 19 |
| Leaf90 | Leaf | Rhizobiales | 21 |
| Leaf361 | Leaf | Rhizobiales | 22 |
| Leaf396 | Leaf | Rhizobiales | 26 |

**Supplementary Table S2:** Comparison of the clades in this study with the clades in Bodilis et al. (2024) and distribution of antiporter per genome and per clade.

| Clade (Bodilis et al., 2024) | Corresponding Clades in This Study | Mean RND Antiporters per genome(All Origins) | Mean RND Antiporters per genome(Plant Origin) | Mean RND Antiporters per genome(Non Plant Origins) | Fold Enrichment (Plant/Non-plant) | Difference (Plant vs Non_Plant Origin) |
| --- | --- | --- | --- | --- | --- | --- |
| *Clade A* | Clade I & Clade II | 2,2 | 4,0 | 2,0 | 2,0 | *** |
| *Clade B* | Clade III | 0,6 | 2,3 | 0,5 | **5,0** | *** |
| *Clade C* | Clade IV | 0,0 | 0,0 | 0,0 | 0,0 | ns |
| *Clade D* | Clade IV | 0,1 | 0,1 | 0,1 | 2,3 | * |
| *Clade E* | Clade IV | 0,2 | 0,5 | 0,2 | 3,1 | *** |
| *Clade F* | Clade V | 0,2 | 0,6 | 0,2 | 3,3 | *** |
| *Clade G* | Clade V | 0,4 | 0,1 | 0,4 | 0,3 | ** |
| *Clade H* | Clade VI | 1,3 | 1,7 | 1,3 | 1,3 | *** |
| *Clade I* | Clade V | 1,5 | 1,5 | 1,5 | 1,0 | ns |
| all clades |  | 6,5 | 10,9 | 6,1 | 1,8 | *** |

\* indicates $p < 0.05$ , \*\* indicates $p < 0.01$ , \*\*\* indicates $p < 0.001$ , while ns indicates no significant differences.

**Supplementary Table S3:** Distribution of RND antiporter homologs across isolate families and plant organs.

| Isolates Order | Isolates Family | Mean antiporter counts per genome |  |  |
| --- | --- | --- | --- | --- |
|  |  | Root Isolates | Leaf Isolates | All Isolates |
| Burkholderiales | Alcaligenaceae | 12.67 ± 0.58 | / | 12.67 ± 0.58 |
| Burkholderiales | Burkholderiaceae | / | 14.00 | 14.00 |
| Burkholderiales | Comamonadaceae | 13.03 ± 2.95 | 10.70 ± 1.42 | 12.44 ± 2.82 |
| Burkholderiales | Oxalobacteraceae | 13.89 ± 1.45 | 15.33 ± 2.89 | 14.25 ± 1.86 |
| Caulobacterales | Caulobacteraceae | 9.29 ± 2.64 | 4.00 ± 0.00 | 8.35 ± 3.16 |
| Cytophagales | Cytophagaceae | / | 16.00 | 16.00 |
| Enterobacterales | Enterobacteriaceae | / | 10.00 ± 0.00 | 10.00 ± 0.00 |
| Flavobacteriales | Flavobacteriaceae | 9.75 ± 1.50 | 7.86 ± 1.57 | 8.55 ± 1.75 |
| Methylophilales | Methylophilaceae | / | 13.67 ± 0.58 | 13.67 ± 0.58 |
| Pseudomonadales | Moraxellaceae | 11.00 | 11.00 | 11.00 ± 0.00 |
| Pseudomonadales | Pseudomonadaceae | 12.00 ± 3.32 | 11.29 ± 2.50 | 11.75 ± 3.01 |
| Rhizobiales | Aurantimonadaceae | / | 7.80 ± 1.30 | 7.80 ± 1.30 |
| Rhizobiales | Bradyrhizobiaceae | 10.40 ± 2.61 | 26.00 | 13.00 ± 6.78 |
| Rhizobiales | Hyphomicrobiaceae | 2.80 ± 1.48 | 5.00 ± 0.00 | 3.43 ± 1.62 |
| Rhizobiales | Methylobacteriaceae | / | 15.28 ± 3.12 | 15.28 ± 3.12 |
| Rhizobiales | Phyllobacteriaceae | 6.75 ± 3.01 | / | 6.75 ± 3.01 |
| Rhizobiales | Rhizobiaceae | 10.56 ± 1.76 | 8.43 ± 0.94 | 9.91 ± 1.84 |
| Sphingobacteriales | Sphingobacteriaceae | / | 7.50 ± 1.29 | 7.50 ± 1.29 |
| Sphingomonadales | Sphingomonadaceae | 6.11 ± 0.78 | 6.41 ± 2.00 | 6.35 ± 1.81 |
| Xanthomonadales | Xanthomonadaceae | 11.69 ± 2.39 | 14.00 ± 3.61 | 12.05 ± 2.63 |

Values are displayed as mean ± sd. "/" indicates absence of data for this Genus/Organ combination.

**Supplementary Table S4.**
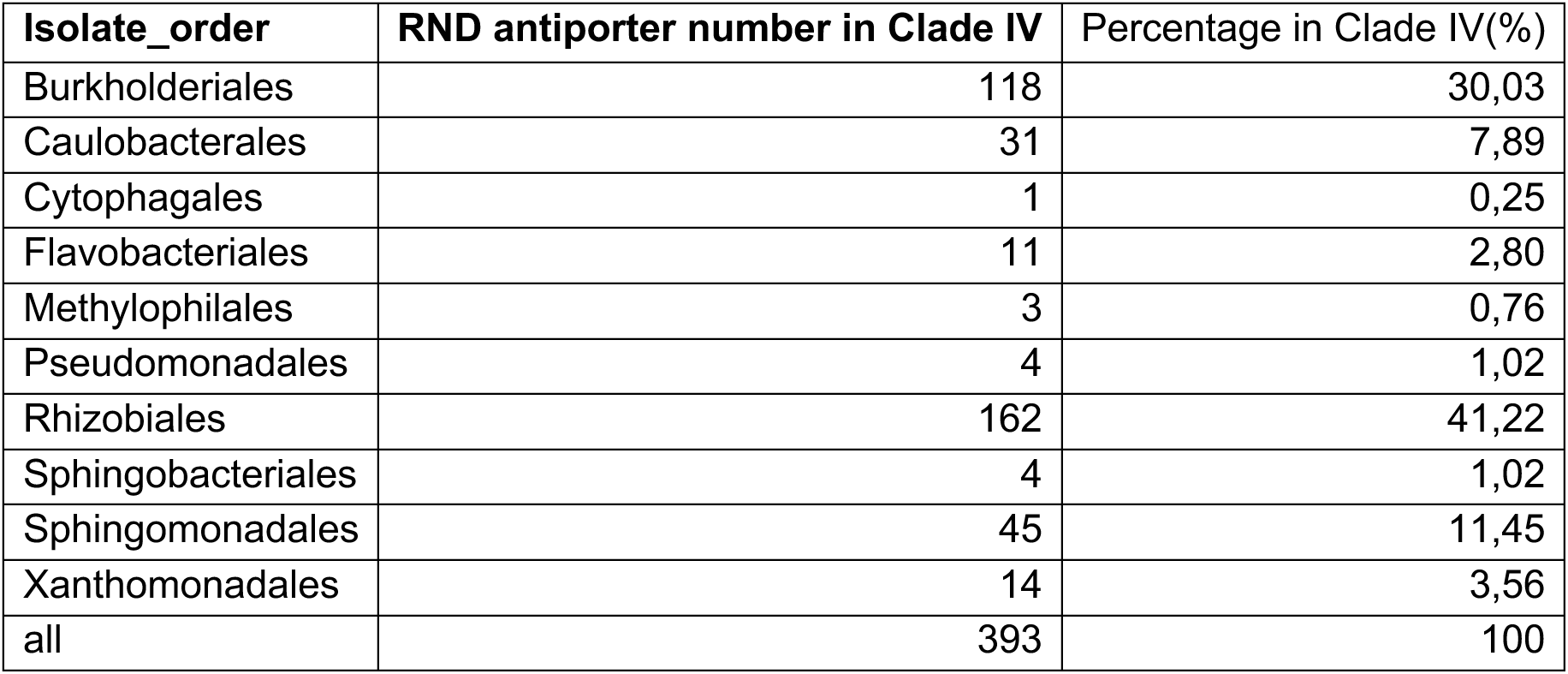
Taxonomic distribution of isolates in Clade IV.

| Isolate_order | RND antiporter number in Clade IV | Percentage in Clade IV(%) |
| --- | --- | --- |
| Burkholderiales | 118 | 30,03 |
| Caulobacterales | 31 | 7,89 |
| Cytophagales | 1 | 0,25 |
| Flavobacteriales | 11 | 2,80 |
| Methylophilales | 3 | 0,76 |
| Pseudomonadales | 4 | 1,02 |
| Rhizobiales | 162 | 41,22 |
| Sphingobacteriales | 4 | 1,02 |
| Sphingomonadales | 45 | 11,45 |
| Xanthomonadales | 14 | 3,56 |
| all | 393 | 100 |

**Supplementary Table S5.**
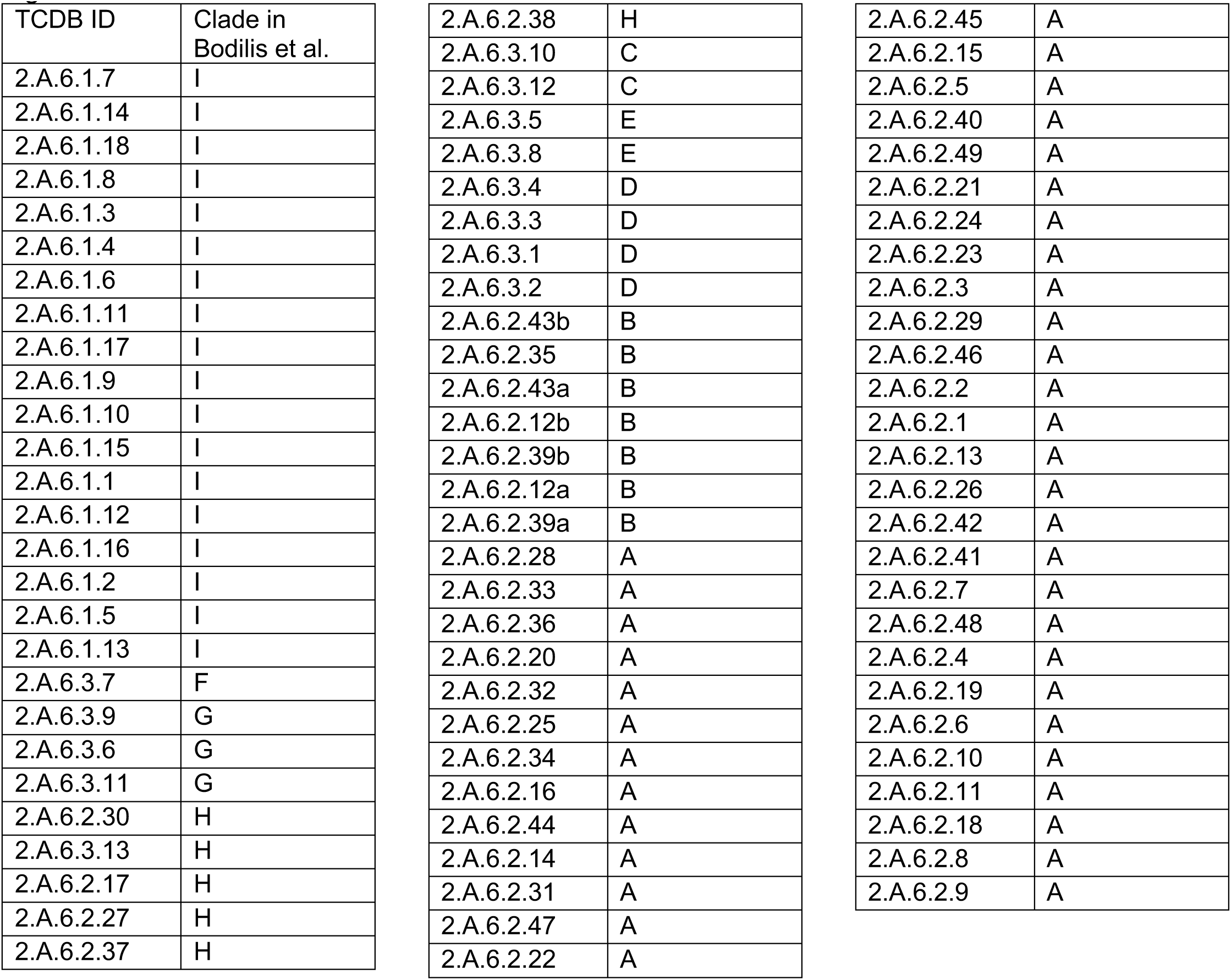
List of transporters from the TCDB database used for Supplementary Figure 2.1a.

| TCDB ID | Clade in Bodilis et al. |  |  |  |  |
| --- | --- | --- | --- | --- | --- |
| 2.A.6.1.7 | I | 2.A.6.2.38 | H | 2.A.6.2.45 | A |
| 2.A.6.1.14 | I | 2.A.6.3.10 | C | 2.A.6.2.15 | A |
| 2.A.6.1.18 | I | 2.A.6.3.12 | C | 2.A.6.2.5 | A |
| 2.A.6.1.8 | I | 2.A.6.3.5 | E | 2.A.6.2.40 | A |
| 2.A.6.1.3 | I | 2.A.6.3.8 | E | 2.A.6.2.49 | A |
| 2.A.6.1.4 | I | 2.A.6.3.4 | D | 2.A.6.2.21 | A |
| 2.A.6.1.6 | I | 2.A.6.3.3 | D | 2.A.6.2.24 | A |
| 2.A.6.1.11 | I | 2.A.6.3.1 | D | 2.A.6.2.23 | A |
| 2.A.6.1.17 | I | 2.A.6.3.2 | D | 2.A.6.2.3 | A |
| 2.A.6.1.9 | I | 2.A.6.2.43b | B | 2.A.6.2.29 | A |
| 2.A.6.1.10 | I | 2.A.6.2.35 | B | 2.A.6.2.46 | A |
| 2.A.6.1.15 | I | 2.A.6.2.43a | B | 2.A.6.2.2 | A |
| 2.A.6.1.1 | I | 2.A.6.2.12b | B | 2.A.6.2.1 | A |
| 2.A.6.1.12 | I | 2.A.6.2.39b | B | 2.A.6.2.13 | A |
| 2.A.6.1.16 | I | 2.A.6.2.12a | B | 2.A.6.2.26 | A |
| 2.A.6.1.2 | I | 2.A.6.2.39a | B | 2.A.6.2.42 | A |
| 2.A.6.1.5 | I | 2.A.6.2.28 | A | 2.A.6.2.41 | A |
| 2.A.6.1.13 | I | 2.A.6.2.33 | A | 2.A.6.2.7 | A |
| 2.A.6.3.7 | F | 2.A.6.2.36 | A | 2.A.6.2.48 | A |
| 2.A.6.3.9 | G | 2.A.6.2.20 | A | 2.A.6.2.4 | A |
| 2.A.6.3.6 | G | 2.A.6.2.32 | A | 2.A.6.2.19 | A |
| 2.A.6.3.11 | G | 2.A.6.2.25 | A | 2.A.6.2.6 | A |
| 2.A.6.2.30 | H | 2.A.6.2.34 | A | 2.A.6.2.10 | A |
| 2.A.6.3.13 | H | 2.A.6.2.16 | A | 2.A.6.2.11 | A |
| 2.A.6.2.17 | H | 2.A.6.2.44 | A | 2.A.6.2.18 | A |
| 2.A.6.2.27 | H | 2.A.6.2.14 | A | 2.A.6.2.8 | A |
| 2.A.6.2.37 | H | 2.A.6.2.31 | A | 2.A.6.2.9 | A |
|  |  | 2.A.6.2.47 | A |  |  |
|  |  | 2.A.6.2.22 | A |  |  |

## References

Abramson, J., Adler, J., Dunger, J., Evans, R., Green, T., Pritzel, A., Ronneberger, O., Willmore, L., Ballard, A.J., Bambrick, J., Bodenstein, S.W., Evans, D.A., Hung, C.-C., O’Neill, M., Reiman, D., Tunyasuvunakool, K., Wu, Z., Žemgulytė, A., Arvaniti, E., Beattie, C., Bertolli, O., Bridgland, A., Cherepanov, A., Congreve, M., Cowen-Rivers, A.I., Cowie, A., Figurnov, M., Fuchs, F.B., Gladman, H., Jain, R., Khan, Y.A., Low, C.M.R., Perlin, K., Potapenko, A., Savy, P., Singh, S., Stecula, A., Thillaisundaram, A., Tong, C., Yakneen, S., Zhong, E.D., Zielinski, M., Žídek, A., Bapst, V., Kohli, P., Jaderberg, M., Hassabis, D., Jumper, J.M., 2024. Accurate structure prediction of biomolecular interactions with AlphaFold 3. Nature 630, 493–500. 10.1038/s41586-024-07487-w

Arnold, B.J., Huang, I.-T., Hanage, W.P., 2022. Horizontal gene transfer and adaptive evolution in bacteria. Nat. Rev. Microbiol. 20, 206–218. 10.1038/s41579-021-00650-4

Baev, N., Endre, G., Petrovics, G., Banfalvi, Z., Kondorosi, A., 1991. Six nodulation genes of nod box locus 4 in Rhizobium meliloti are involved in nodulation signal production: nodM codes for d-glucosamine synthetase. Mol. Gen. Genet. MGG 228, 113–124. 10.1007/BF00282455

Bai, Y., Müller, D.B., Srinivas, G., Garrido-Oter, R., Potthoff, E., Rott, M., Dombrowski, N., Münch, P.C., Spaepen, S., Remus-Emsermann, M., Hüttel, B., McHardy, A.C., Vorholt, J.A., Schulze-Lefert, P., 2015. Functional overlap of the Arabidopsis leaf and root microbiota. Nature 528, 364–369. 10.1038/nature16192

Bailey, T.L., 2021. STREME: accurate and versatile sequence motif discovery. Bioinformatics 37, 2834–2840. 10.1093/bioinformatics/btab203

Barillot, C.D.C., Sarde, C.-O., Bert, V., Tarnaud, E., Cochet, N., 2013. A standardized method for the sampling of rhizosphere and rhizoplan soil bacteria associated to a herbaceous root system. Ann. Microbiol. 63, 471–476. 10.1007/s13213-012-0491-y

Barnett, M.J., Fisher, R.F., Jones, T., Komp, C., Abola, A.P., Barloy-Hubler, F., Bowser, L., Capela, D., Galibert, F., Gouzy, J., Gurjal, M., Hong, A., Huizar, L., Hyman, R.W., Kahn, D., Kahn, M.L., Kalman, S., Keating, D.H., Palm, C., Peck, M.C., Surzycki, R., Wells, D.H., Yeh, K.-C., Davis, R.W., Federspiel, N.A., Long, S.R., 2001. Nucleotide sequence and predicted functions of the entire *Sinorhizobium meliloti* pSymA megaplasmid. Proc. Natl. Acad. Sci. 98, 9883–9888. 10.1073/pnas.161294798

Black, M., Moolhuijzen, P., Chapman, B., Barrero, R., Howieson, J., Hungria, M., Bellgard, M., 2012. The Genetics of Symbiotic Nitrogen Fixation: Comparative Genomics of 14 Rhizobia Strains by Resolution of Protein Clusters. Genes 3, 138–166. 10.3390/genes3010138

Blair, J.M., Richmond, G.E., Piddock, L.J., 2014. Multidrug Efflux Pumps in Gram-Negative Bacteria and Their Role in Antibiotic Resistance. Future Microbiol. 9, 1165–1177. 10.2217/fmb.14.66

Blanco, P., Hernando-Amado, S., Reales-Calderon, J.A., Corona, F., Lira, F., Alcalde-Rico, M., Bernardini, A., Sanchez, M.B., Martinez, J.L., 2016. Bacterial Multidrug Efflux Pumps: Much More Than Antibiotic Resistance Determinants. Microorganisms 4, 14. 10.3390/microorganisms4010014

Bodilis, J., Simenel, O., Michalet, S., Brothier, E., Meyer, T., Favre-Bonté, S., Nazaret, S., 2024. HME, NFE, and HAE-1 efflux pumps in Gram-negative bacteria: a comprehensive phylogenetic and ecological approach. ISME Commun. 4, ycad018. 10.1093/ismeco/ycad018

Bulgarelli, D., Schlaeppi, K., Spaepen, S., van Themaat, E.V.L., Schulze-Lefert, P., 2013. Structure and Functions of the Bacterial Microbiota of Plants. Annu. Rev. Plant Biol. 64, 807–838. 10.1146/annurev-arplant-050312-120106

Camacho, C., Coulouris, G., Avagyan, V., Ma, N., Papadopoulos, J., Bealer, K., Madden, T.L., 2009. BLAST+: architecture and applications. BMC Bioinformatics 10, 421. 10.1186/1471-2105-10-421

Canarini, A., Kaiser, C., Merchant, A., Richter, A., Wanek, W., 2019. Root Exudation of Primary Metabolites: Mechanisms and Their Roles in Plant Responses to Environmental Stimuli. Front. Plant Sci. 10. 10.3389/fpls.2019.00157

Chopra, I., Roberts, M., 2001. Tetracycline Antibiotics: Mode of Action, Applications, Molecular Biology, and Epidemiology of Bacterial Resistance. Microbiol. Mol. Biol. Rev. 65, 232–260. 10.1128/mmbr.65.2.232-260.2001

Daury, L., Orange, F., Taveau, J.-C., Verchère, A., Monlezun, L., Gounou, C., Marreddy, R.K.R., Picard, M., Broutin, I., Pos, K.M., Lambert, O., 2016. Tripartite assembly of RND multidrug efflux pumps. Nat. Commun. 7, 10731. 10.1038/ncomms10731

Dénarié, J., Debellé, F., Promé, J.C., 1996. Rhizobium lipo-chitooligosaccharide nodulation factors: signaling molecules mediating recognition and morphogenesis. Annu. Rev. Biochem. 65, 503–535. 10.1146/annurev.bi.65.070196.002443

Drigo, B., Pijl, A.S., Duyts, H., Kielak, A.M., Gamper, H.A., Houtekamer, M.J., Boschker, H.T.S., Bodelier, P.L.E., Whiteley, A.S., Veen, J.A.V., Kowalchuk, G.A., 2010. Shifting carbon flow from roots into associated microbial communities in response to elevated atmospheric CO_2_. Proc. Natl. Acad. Sci. 107, 10938–10942. 10.1073/pnas.0912421107

Du, D., Wang-Kan, X., Neuberger, A., van Veen, H.W., Pos, K.M., Piddock, L.J.V., Luisi, B.F., 2018. Multidrug efflux pumps: structure, function and regulation. Nat. Rev. Microbiol. 16, 523–539. 10.1038/s41579-018-0048-6

Durán, P., Thiergart, T., Garrido-Oter, R., Agler, M., Kemen, E., Schulze-Lefert, P., Hacquard, S., 2018. Microbial Interkingdom Interactions in Roots Promote Arabidopsis Survival. Cell 175, 973–983.e14. 10.1016/j.cell.2018.10.020

Fanelli, G., Pasqua, M., Prosseda, G., Grossi, M., Colonna, B., 2023. AcrAB efflux pump impacts on the survival of adherent-invasive Escherichia coli strain LF82 inside macrophages. Sci. Rep. 13, 2692. 10.1038/s41598-023-29817-0

Finkel, O.M., Salas-González, I., Castrillo, G., Conway, J.M., Law, T.F., Teixeira, P.J.P.L., Wilson, E.D., Fitzpatrick, C.R., Jones, C.D., Dangl, J.L., 2020. A single bacterial genus maintains root growth in a complex microbiome. Nature 587, 103–108. 10.1038/s41586-020-2778-7

Fitzpatrick, C.R., Salas-González, I., Conway, J.M., Finkel, O.M., Gilbert, S., Russ, D., Teixeira, P.J.P.L., Dangl, J.L., 2020. The Plant Microbiome: From Ecology to Reductionism and Beyond. Annu. Rev. Microbiol. 74, 81–100. 10.1146/annurev-micro-022620-014327

Grant, B.J., Rodrigues, A.P.C., ElSawy, K.M., McCammon, J.A., Caves, L.S.D., 2006. Bio3d: an R package for the comparative analysis of protein structures. Bioinformatics 22, 2695–2696. 10.1093/bioinformatics/btl461

Hacquard, S., Garrido-Oter, R., González, A., Spaepen, S., Ackermann, G., Lebeis, S., McHardy, A.C., Dangl, J.L., Knight, R., Ley, R., Schulze-Lefert, P., 2015. Microbiota and Host Nutrition across Plant and Animal Kingdoms. Cell Host Microbe 17, 603–616. 10.1016/j.chom.2015.04.009

Harbort, C.J., Hashimoto, M., Inoue, H., Schulze-Lefert, P., 2020. A gnotobiotic growth assay for Arabidopsis root microbiota reconstitution under iron limitation. STAR Protoc. 1, 100226. 10.1016/j.xpro.2020.100226

Hoang, D.T., Chernomor, O., von Haeseler, A., Minh, B.Q., Vinh, L.S., 2018. UFBoot2: Improving the Ultrafast Bootstrap Approximation. Mol. Biol. Evol. 35, 518–522. 10.1093/molbev/msx281

Kalyaanamoorthy, S., Minh, B.Q., Wong, T.K.F., Von Haeseler, A., Jermiin, L.S., 2017. ModelFinder: fast model selection for accurate phylogenetic estimates. Nat. Methods 14, 587–589. 10.1038/nmeth.4285

Katoh, K., Misawa, K., Kuma, K., Miyata, T., 2002. MAFFT: a novel method for rapid multiple sequence alignment based on fast Fourier transform. Nucleic Acids Res. 30, 3059–3066. 10.1093/nar/gkf436

Kim, J.-Y., Loo, E.P.-I., Pang, T.Y., Lercher, M., Frommer, W.B., Wudick, M.M., 2021. Cellular export of sugars and amino acids: role in feeding other cells and organisms. Plant Physiol. 187, 1893–1914. 10.1093/plphys/kiab228

Kobylka, J., Kuth, M.S., Müller, R.T., Geertsma, E.R., Pos, K.M., 2020. AcrB: a mean, keen, drug efflux machine. Ann. N. Y. Acad. Sci. 1459, 38–68. 10.1111/nyas.14239

Lian, B., Prithiviraj, B., Souleimanov, A., Smith, D.L., 2001. Evidence for the production of chemical compounds analogous to nod factor by the silicate bacterium Bacillus circulans GY92. Microbiol. Res. 156, 289–292. 10.1078/0944-5013-00107

Martinez, J.L., Sánchez, M.B., Martínez-Solano, L., Hernandez, A., Garmendia, L., Fajardo, A., Alvarez-Ortega, C., 2009. Functional role of bacterial multidrug efflux pumps in microbial natural ecosystems. FEMS Microbiol. Rev. 33, 430–449. 10.1111/j.1574-6976.2008.00157.x

Meng, E.C., Goddard, T.D., Pettersen, E.F., Couch, G.S., Pearson, Z.J., Morris, J.H., Ferrin, T.E., 2023. UCSF ChimeraX: Tools for structure building and analysis. Protein Sci. 32, e4792. 10.1002/pro.4792

Mercier, J., Lindow, S.E., 2000. Role of Leaf Surface Sugars in Colonization of Plants by Bacterial Epiphytes. Appl. Environ. Microbiol. 66, 369–374. 10.1128/aem.66.1.369-374.2000

Minh, B.Q., Schmidt, H.A., Chernomor, O., Schrempf, D., Woodhams, M.D., von Haeseler, A., Lanfear, R., 2020. IQ-TREE 2: New Models and Efficient Methods for Phylogenetic Inference in the Genomic Era. Mol. Biol. Evol. 37, 1530–1534. 10.1093/molbev/msaa015

Murakami, S., Nakashima, R., Yamashita, E., Yamaguchi, A., 2002. Crystal structure of bacterial multidrug efflux transporter AcrB. Nature 419, 587–593. 10.1038/nature01050

Murrell, B., Moola, S., Mabona, A., Weighill, T., Sheward, D., Kosakovsky Pond, S.L., Scheffler, K., 2013. FUBAR: A Fast, Unconstrained Bayesian AppRoximation for Inferring Selection. Mol. Biol. Evol. 30, 1196–1205. 10.1093/molbev/mst030

Oberai, A., Joh, N.H., Pettit, F.K., Bowie, J.U., 2009. Structural imperatives impose diverse evolutionary constraints on helical membrane proteins. Proc. Natl. Acad. Sci. U. S. A. 106, 17747–17750. 10.1073/pnas.0906390106

Parniske, M., 1991. Flavonoide als Signal- und Abwehrstoffe in der Glycine/Bradyrhizobium Symbiose. Marburg.

Parniske, M., Ahlborn, B., Werner, D., 1991. Isoflavonoid-inducible resistance to the phytoalexin glyceollin in soybean rhizobia. J. Bacteriol. 173, 3432–3439. 10.1128/jb.173.11.3432-3439.1991

Piddock, L.J.V., 2006. Multidrug-resistance efflux pumps - not just for resistance. Nat. Rev. Microbiol. 4, 629–636. 10.1038/nrmicro1464

Pond, S.L.K., Frost, S.D.W., Muse, S.V., 2005. HyPhy: hypothesis testing using phylogenies. Bioinformatics 21, 676–679. 10.1093/bioinformatics/bti079

Rush, T.A., Puech-Pagès, V., Bascaules, A., Jargeat, P., Maillet, F., Haouy, A., Maës, A.Q., Carriel, C.C., Khokhani, D., Keller-Pearson, M., Tannous, J., Cope, K.R., Garcia, K., Maeda, J., Johnson, C., Kleven, B., Choudhury, Q.J., Labbé, J., Swift, C., O’Malley, M.A., Bok, J.W., Cottaz, S., Fort, S., Poinsot, V., Sussman, M.R., Lefort, C., Nett, J., Keller, N.P., Bécard, G., Ané, J.-M., 2020. Lipo-chitooligosaccharides as regulatory signals of fungal growth and development. Nat. Commun. 11, 3897. 10.1038/s41467-020-17615-5

Russ, D., Fitzpatrick, C.R., Saha, C., Law, T.F., Jones, C.D., Kliebenstein, D.J., Dangl, J.L., 2025. An effluent pump family distributed across plant commensal bacteria conditions host- and organ-specific glucosinolate detoxification. Nat. Commun. 16, 5699. 10.1038/s41467-025-61266-3

Saier Jr, M.H., Tam, R., Reizer, A., Reizer, J., 1994. Two novel families of bacterial membrane proteins concerned with nodulation, cell division and transport. Mol. Microbiol. 11, 841–847. 10.1111/j.1365-2958.1994.tb00362.x

Saier, M.H., Tran, C.V., Barabote, R.D., 2006. TCDB: the Transporter Classification Database for membrane transport protein analyses and information. Nucleic Acids Res. 34, D181–186. 10.1093/nar/gkj001

Santus, L., Espinosa-Carrasco, J., Rauschning, L., alessiovignoli, Trujnara, I., bot, nf-core, Klaps, J., 2024. nf-core/multiplesequencealign: nf-core/multiplesequencealign v1.0.0 - Somorrostro. 10.5281/zenodo.13889387

Sasse, J., Martinoia, E., Northen, T., 2018. Feed Your Friends: Do Plant Exudates Shape the Root Microbiome? Trends Plant Sci. 23, 25–41. 10.1016/j.tplants.2017.09.003

Schäfer, M., Pacheco, A.R., Künzler, R., Bortfeld-Miller, M., Field, C.M., Vayena, E., Hatzimanikatis, V., Vorholt, J.A., 2023. Metabolic interaction models recapitulate leaf microbiota ecology. Science 381, eadf5121. 10.1126/science.adf5121

Schandry, N., Jandrasits, K., Garrido-Oter, R., Becker, C., 2021. Plant-derived benzoxazinoids act as antibiotics and shape bacterial communities. 10.1101/2021.01.12.425818

Schwengers, O., Jelonek, L., Dieckmann, M.A., Beyvers, S., Blom, J., Goesmann, A., 2021. Bakta: rapid and standardized annotation of bacterial genomes via alignment-free sequence identification. Microb. Genomics 7, 000685. 10.1099/mgen.0.000685

Tam, H.-K., Foong, W.E., Oswald, C., Herrmann, A., Zeng, H., Pos, K.M., 2021. Allosteric drug transport mechanism of multidrug transporter AcrB. Nat. Commun. 12, 3889. 10.1038/s41467-021-24151-3

Terán, W., Felipe, A., Segura, A., Rojas, A., Ramos, J.-L., Gallegos, M.-T., 2003. Antibiotic-Dependent Induction of Pseudomonas putida DOT-T1E TtgABC Efflux Pump Is Mediated by the Drug Binding Repressor TtgR. Antimicrob. Agents Chemother. 47, 3067–3072. 10.1128/aac.47.10.3067-3072.2003

Thoenen, L., Giroud, C., Kreuzer, M., Waelchli, J., Gfeller, V., Deslandes-Hérold, G., Mateo, P., Robert, C.A.M., Ahrens, C.H., Rubio-Somoza, I., Bruggmann, R., Erb, M., Schlaeppi, K., 2023. Bacterial tolerance to host-exuded specialized metabolites structures the maize root microbiome. Proc. Natl. Acad. Sci. 120, e2310134120. 10.1073/pnas.2310134120

Truchet, G., Roche, P., Lerouge, P., Vasse, J., Camut, S., de Billy, F., Promé, J.-C., Dénarié, J., 1991. Sulphated lipo-oligosaccharide signals of Rhizobium meliloti elicit root nodule organogenesis in alfalfa. Nature 351, 670–673. 10.1038/351670a0

Voges, M.J.E.E.E., Bai, Y., Schulze-Lefert, P., Sattely, E.S., 2019. Plant-derived coumarins shape the composition of an *Arabidopsis* synthetic root microbiome. Proc. Natl. Acad. Sci. 116, 12558–12565. 10.1073/pnas.1820691116

Vorholt, J.A., 2012. Microbial life in the phyllosphere. Nat. Rev. Microbiol. 10, 828–840. 10.1038/nrmicro2910

Wang-Kan, X., Blair, J.M.A., Chirullo, B., Betts, J., La Ragione, R.M., Ivens, A., Ricci, V., Opperman, T.J., Piddock, L.J.V., 2017. Lack of AcrB Efflux Function Confers Loss of Virulence on *Salmonella enterica* Serovar Typhimurium. mBio 8, e00968–17. 10.1128/mBio.00968-17

Wilhelm, J., Pos, K.M., 2024. Molecular insights into the determinants of substrate specificity and efflux inhibition of the RND efflux pumps AcrB and AdeB. Microbiology 170, 001438. 10.1099/mic.0.001438

Xu, S., Dai, Z., Guo, P., Fu, X., Liu, S., Zhou, L., Tang, W., Feng, T., Chen, M., Zhan, L., Wu, T., Hu, E., Jiang, Y., Bo, X., Yu, G., 2021. ggtreeExtra: Compact Visualization of Richly Annotated Phylogenetic Data. Mol. Biol. Evol. 38, 4039–4042. 10.1093/molbev/msab166

Yu, E.W., McDermott, G., Zgurskaya, H.I., Nikaido, H., Koshland, D.E., 2003. Structural basis of multiple drug-binding capacity of the AcrB multidrug efflux pump. Science 300, 976–980. 10.1126/science.1083137

Yu, G., 2020. Using ggtree to Visualize Data on Tree-Like Structures. Curr. Protoc. Bioinforma. 69. 10.1002/cpbi.96

Zahedi Bialvaei, A., Rahbar, M., Hamidi-Farahani, R., Asgari, A., Esmailkhani, A., Mardani Dashti, Y., Soleiman-Meigooni, S., 2021. Expression of RND efflux pumps mediated antibiotic resistance in Pseudomonas aeruginosa clinical strains. Microb. Pathog. 153, 104789. 10.1016/j.micpath.2021.104789

